# Deciphering the nucleotide driven kinetic and oligomeric dynamics in IMPDH regulation

**DOI:** 10.64898/2026.07.01.735766

**Authors:** Nour Ayoub, Bertrand Raynal, Muriel Gelin, Gilles Labesse, Ahmed Haouz, Hélène Munier-Lehmann

**Affiliations:** Université Paris Cité, INSERM, Health & Functional Exposomics - HealthFex, U1124, Institut Pasteur, Structural Biology and Chemistry Department, F-75006 Paris, France; Institut Pasteur, Université Paris Cité, CNRS UMR3528, Plate-Forme de Biophysique Moléculaire, F-75015 Paris, France; Centre de Biologie Structurale, Université de Montpellier, INSERM, CNRS, F-34090 Montpellier, France; Institut Pasteur, Université Paris Cité, CNRS UMR3528, Plate-Forme de Cristallographie, F-75015 Paris, France

**Keywords:** de novo purine biosynthesis, IMPDH, Burkholderia thailandensis, Escherichia coli, Pseudomonas aeruginosa, integrative structural biology

## Abstract

Inosine monophosphate dehydrogenase (IMPDH) controls guanine nucleotide biosynthesis. Here, using biochemical and integrative structural biology approaches, we characterize the class II bacterial IMPDH from *Burkholderia thailandensis*. We demonstrate that MgGTP acts as a direct allosteric inhibitor independently of MgATP, promoting tetramer-to-octamer assembly via the Bateman domain. When both nucleotides are present, the enzyme exhibits a biphasic response: low MgGTP concentrations enhance activity, whereas higher concentrations restore inhibition. Structural analyses reveal distinct octameric conformations and capture a pre-catalytic Michaelis complex with substrates and effectors bound. These findings uncover a regulatory mechanism where the Bateman domain integrates opposing nucleotide signals to dynamically control guanine nucleotide biosynthesis, highlighting IMPDH as a potential antimicrobial target in pathogenic *Burkholderia* species.

## Introduction

Guanine nucleotide homeostasis is essential for cellular physiology, as the balance between adenosine- and guanosine-derived nucleotides underpins processes ranging from energy metabolism to nucleic acid synthesis, translation, and signal transduction. A central regulator of this balance is inosine monophosphate dehydrogenase (IMPDH), which catalyzes the rate-limiting step in the *de novo* biosynthesis of guanine nucleotides by converting inosine monophosphate (IMP) into xanthosine monophosphate (XMP) with concomitant reduction of NAD⁺ to NADH. By controlling metabolic flux toward guanine nucleotide synthesis, IMPDH plays a pivotal role in maintaining the balance between intracellular ATP and GTP pools, thereby ensuring proper nucleotide homeostasis and supporting essential cellular processes such as growth and proliferation^1^.

IMPDHs are widely conserved among bacteria, archaea, and eukaryotes, yet they display remarkable diversity in their catalytic and regulatory properties^1^. All canonical IMPDHs share a common architecture consisting of a regulatory Bateman domain^2^ (BD; also called CBS domains or CBS modules) nested within a loop of the catalytic (β/α)8 barrel-domain (CD), thus dividing the catalytic domain into two fragments, commonly referred to as CD1 and CD2. The Bateman domain acts as a major allosteric regulatory module that binds adenine and guanine nucleotides and modulates the enzyme’s activity in response to metabolic signals. Comparisons of quaternary structures and catalytic properties have revealed the existence of two functional classes of IMPDHs that differ in their modes of allosteric regulation and conformational dynamics^3^. These classes reflect evolutionary adaptations that fine-tune enzyme activity to specific cellular contexts. Notably, characterization of chimeric IMPDHs in which Bateman domains were swapped between class I and class II enzymes, demonstrated that this domain serves as the modular determinant of functional behavior. The Bateman domain dictates differential allosteric responses and oligomeric behaviors observed in the two classes, highlighting its central role in controlling IMPDH regulation^3^.

Despite extensive characterization, the molecular mechanisms underlying nucleotide-dependent regulation in class II IMPDHs remain incompletely understood. In several bacterial IMPDHs, a variety of nucleotides beyond MgATP have been shown to bind the Bateman domain and modulate enzyme activity through distinct regulatory mechanisms^1^. However, whether such ligands can differentially regulate the catalytic activity and oligomeric state of class II IMPDHs remains unclear. Exploring how nucleotides other than MgATP influence these properties is therefore essential for understanding how IMPDH integrates metabolic signals to dynamically adjust guanine nucleotide biosynthesis to cellular demands.

To address these questions, we focused our investigation on class II bacterial IMPDHs and selected the enzyme from *Burkholderia thailandensis* (IMPDHbt) as a representative. This enzyme was chosen as our model system for two reasons. First, it provides a valuable system for elucidating the fundamental mechanisms of class II IMPDH regulation. Second, IMPDH remains a completely unexplored therapeutic target within the *Burkholderia* genus. *B. thailandensis* is a close relative of *B. pseudomallei*, the causative agent of melioidosis^4, 5^ that is classified as a Tier 1 Select Agent due to the stringent biosafety and security measures required for pathogens with high potential public health impact, as well as of the members of the *B. cepacia* complex (Bcc)^5, 6^, which cause fatal disseminated infections in immunocompromised hosts. Thus, *B. thailandensis* provides an accessible model for investigating IMPDH regulation in those pathogens. The ability of these pathogens to survive in both intracellularly and extracellular environments during systemic infection makes their purine nucleotide metabolism a potentially attractive yet unexplored target for antimicrobial intervention^7^. Understanding how the Bateman domain mediates allosteric communication with the catalytic core is therefore essential for deciphering the mechanistic basis governing IMPDH regulation in these clinically relevant pathogens.

Here we investigate the mechanisms governing nucleotide-dependent regulation of IMPDHbt as well as the Bateman-domain deleted variant (IMPDHbt ΔDB). Using a combination of biochemical and integrative structural biology approaches, we show that guanine nucleotides differentially modulate the catalytic activity and assembly state of IMPDHbt. Our results reveal how ligand binding to the Bateman domain is coupled to structural changes that propagate to the catalytic core, uncovering previously unrecognized regulatory mechanisms that control IMPDH function in class II bacterial enzymes. These findings provide new insights into the allosteric regulation of guanine nucleotide biosynthesis and highlight IMPDH as a potential metabolic vulnerability in pathogenic *Burkholderia* species.

## Results

### Nucleotide-mediated regulation of class II bacterial IMPDHs

All recombinant IMPDHs were overexpressed in *E*. *coli* and purified using a standard two-step procedure (see Methods) as previously described^3, 8, 9^. *In vitro* enzymatic assays were performed by monitoring NADH production at 340 nm under non-saturating substrate conditions (0.2 mM IMP, 0.4 mM NAD⁺, and 3 mM MgCl₂) in the absence or in the presence of various nucleotides (1 mM), including guanosine, GMP, GDP, GTP, cGMP, cAMP, cCMP and XMP. Among the nucleotides tested, GMP, XMP, GDP and GTP significantly inhibited the IMPDHbt catalytic activity (between 40% and 83%), whereas the other compounds did not exert any notable effect (Fig. 1a). To evaluate the contribution of magnesium in the observed inhibition, the activity of IMPDHbt was monitored with or without equimolar MgCl₂ for GMP and GTP. While GMP-mediated inhibition was unaffected (Fig. 1b), GTP inhibition was enhanced in the presence of MgCl₂ (Fig. 1c). These observations prompted the selection of GMP and MgGTP for further analyses. As expected, and already depicted for other IMPDHs^10^, GMP was shown to be a competitive inhibitor of IMP (Supplementary Fig. 1). Secondary plots of K_m_^app^ as a function of GMP concentration led to the determination of the inhibition constant of IMPDHbt for GMP (Kᵢ^GMP^ = 0.08 ± 0.02 mM). Regarding MgGTP, we tested the impact of MgATP on the observed MgGTP-mediated inhibition of the IMPDHbt catalytic activity (Fig.1d-e). A biphasic response to MgGTP was measured in the presence of 2 mM MgATP: sub-millimolar concentrations of MgGTP (<0.2 mM) enhanced activity, while higher concentrations were inhibitory (Fig. 1d). Conversely, varying MgATP in the presence of 2 mM MgGTP highlighted a reversal of inhibition only above 0.3 mM MgATP, suggesting a complex mechanism involving both nucleotides (Fig. 1e).

**Figure 1.**
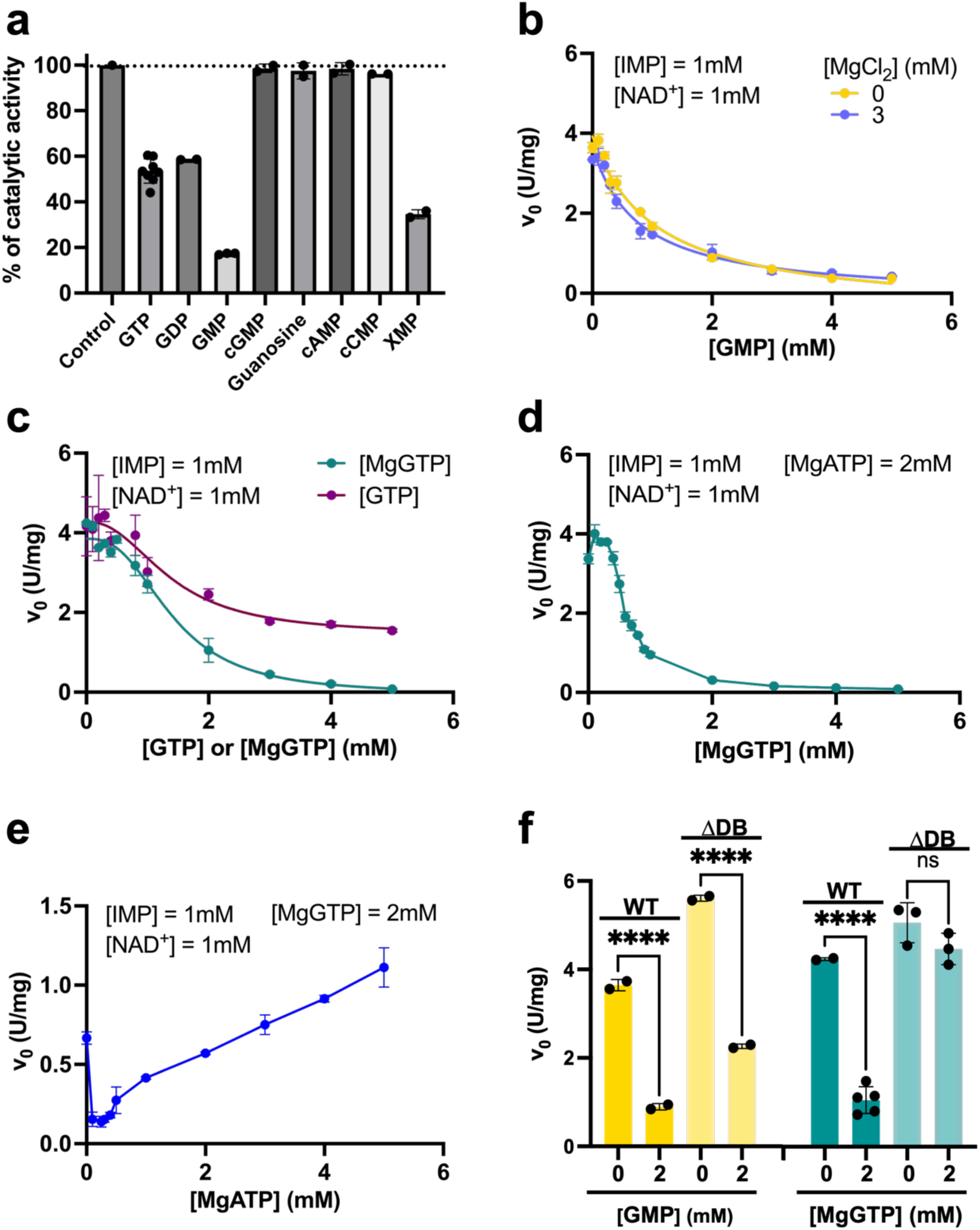
Kinetic analysis of IMPDHbt in the presence of purine nucleotides. **a** Effect of various nucleotides at a concentration of 1 mM. The catalytic activity measured in the presence of 0.2 mM IMP, 0.4 NAD^+^ and 3 mM MgCl_2_ was taken as the control and defined as 100%. **b** Variation of the initial velocity, fitted with the “[inhibitor] versus response” equation (GraphPad), as a function of GMP in the absence or in the presence of MgCl_2_. **c** Same as in **b** for GTP. **d** Variation of the initial velocity of IMPDHbt as a function of MgGTP concentration in the presence of 2 mM MgATP. **e** Variation of the initial velocity of IMPDHbt as a function of MgATP concentration in the presence of 1 mM IMP, 1 mM NAD^+^ and 2 mM MgGTP. **f** Histogram showing the initial velocity of the wild-type and ΔDB IMPDHbt variant in the absence or in the presence of 2 mM GMP or MgGTP, at fixed concentrations of 1 mM IMP and 1 mM NAD⁺. Data were obtained from at least three independent measurements. Each data point represents a mean ± standard deviation in error bars.

To further explore the role of the Bateman domain in nucleotide sensing, enzymatic activity was evaluated using a variant of IMPDHbt deleted of its Bateman domain (IMPDHbt ΔDB). Despite retaining GMP sensitivity, this variant lost its response to MgGTP (Fig. 1f), strongly indicating that MgGTP binds specifically to the Bateman domain.

The inhibition by MgGTP alone observed on IMPDHbt led us to investigate its impact on the catalytic activity of other bacterial IMPDHs. Thus, we performed similar kinetic experiments on another class II IMPDH (i.e. *Escherichia coli* IMPDH (IMPDHec)) and on a representative of class I IMPDH (i.e. *Pseudomonas aeruginosa* IMPDH (IMPDHpa)). For both IMPDHs (Supplementary Fig. 2) MgGTP is an inhibitor with an I_50_ in the millimolar range as for IMPDHbt (Supplementary Table 1).

Enzymatic activity was then assessed with varying concentrations of IMP or NAD⁺ in the absence or the presence of 2 mM MgGTP (Fig. 2). Among the three bacterial IMPDHs (IMPDHbt, IMPDHec, IMPDHpa) tested, both class II enzymes displayed hyperbolic kinetics, consistent with a Michaelis-Menten model, while for IMPDHpa, its activity profile deviated significantly (Fig. 2f)) as previously reported^3, 8, 9^. MgGTP increases both the Hill number and the apparent K_0.5_ for IMP, while decreasing the reaction rate (Table 1). On the other hand, the affinity for the second substrate NAD^+^ was enhanced. For the two class II IMPDHs (Table 1), MgGTP also impacted the reaction rates and in some cases the substrate affinity. Overall, for the three bacterial IMPDHs, MgGTP acted as a negative effector, with drastic impact on the kinetic parameters (a reduction by up to 2- and 6.2-fold for the V_m_ or the affinity for IMP, respectively).

**Figure 2.**
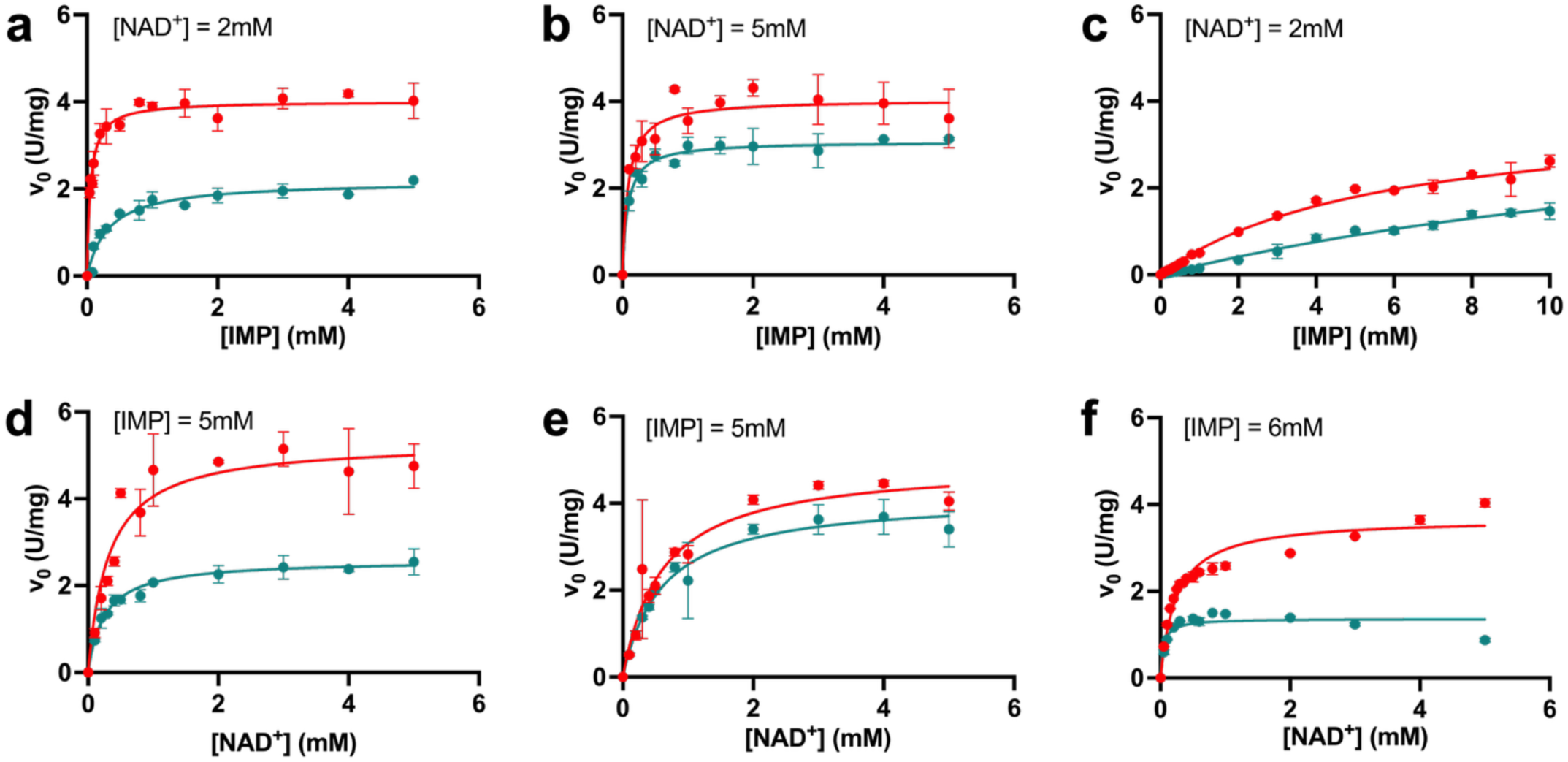
IMPDHbt, IMPDHec and IMPDHpa are inhibited by MgGTP in a MgATP-independent manner. Initial velocity curves as a function of IMP concentration for IMPDHbt (**a**), IMPDHec (**b**), and IMPDHpa (**c**), in the absence (red) or in the presence (green) of 2 mM MgGTP (3 mM for IMPDHpa). Fixed NAD⁺ concentrations were 2 mM for IMPDHbt and IMPDHpa, and 5 mM for IMPDHec. Measurements (performed in at least triplicate) were fitted using the Michaelis-Menten equation, except for IMPDHpa, which was fitted using the Hill equation. The same experiments were performed at a fixed IMP concentration (5 mM for IMPDHbt and IMPDHec, 6 mM for IMPDHpa) and with variable NAD^+^ concentration for IMPDHbt (**d**), IMPDHec (**e**) and IMPDHpa (**f**) in the absence (red) or in the presence (green) of 2 mM MgGTP (3 mM for IMPDHpa). Measurements (performed in at least triplicate) were fitted using the Michaelis–Menten equation. Each data point represents a mean ± standard deviation in error bars. All the calculated parameters are displayed in **Table 1**.

**Table 1.** Kinetic parameters of IMPDHbt, IMPDHec and IMPDHpa, with IMP or NAD^+^ as variable substrate. Reaction rates were determined at a constant concentration of NAD^+^ (2 mM for IMPDHbt and IMPDHpa; 5 mM for IMPDHec) or at a constant concentration of IMP (2 mM for IMPDHbt and IMPDHpa; 5 mM for IMPDHec) and in the absence or in the presence of MgGTP (5 mM for IMPDHbt and IMPDHec; 6 mM for IMPDHpa), and fitted according to the Michaelis-Menten equation (IMPDHbt, IMPDHec and IMPDHpa for NAD^+^ as a variable substrate) or the Hill equation (IMPDHpa for IMP as a variable substrate).

| Without MgGTP |  |  |  | With MgGTP |  |  |
| --- | --- | --- | --- | --- | --- | --- |
| [IMP] variable | V<sub>m</sub> (U/mg) | K<sub>m</sub><sup>IMP</sup> or K<sub>0.5</sub><sup>IMP</sup> (μM) | n<sub>H</sub> | V<sub>m</sub> (U/mg) | K<sub>m</sub><sup>IMP</sup> or K<sub>0.5</sub><sup>IMP</sup> (μM) | n<sub>H</sub> |
| IMPDHbt | 4.01 ± 0.06 | 53 ± 5 | - | 2.18 ± 0.08 | 330 ± 46 | - |
| IMPDHec | 4.04 ± 0.13 | 87 ± 19 | - | 3.07 ± 0.07 | 82 ± 14 | - |
| IMPDHpa | 3.11 ± 0.30 | 3648 ± 726 | 1.20 ± 0.11 | 2.36 ± 0.38 | 6736 ± 1709 | 1.36 ± 0.16 |
| [NAD<sup>+</sup>] variable | V<sub>m</sub> (U/mg) | K<sub>m</sub><sup>NAD<sup>+</sup></sup> (μM) |  | V<sub>m</sub> (U/mg) | K<sub>m</sub><sup>NAD<sup>+</sup></sup> (μM) |  |
| IMPDHbt | 5.31 ± 0.24 | 311 ± 61 |  | 2.59 ± 0.07 | 250 ± 29 |  |
| IMPDHec | 4.92 ± 0.26 | 599 ± 99 |  | 4.16 ± 0.16 | 604 ± 80 |  |
| IMPDHpa | 3.68 ± 0.10 | 242 ± 25 |  | 1.37 ± 0.06 | 44 ± 14 |  |

### Regulation of IMPDHbt oligomeric state

We have previously shown that the oligomeric states of class II IMPDHs were regulated by different nucleotides bound to the catalytic or the Bateman domains (NAD^+^ and MgATP, respectively)^8^. To gain insight into the structural basis of IMPDHbt regulation by GMP and MgGTP, its oligomeric state was investigated using analytical ultracentrifugation (AUC) and mass photometry (MP). Both AUC and MP analyses (Fig. 3) confirmed the existence of a tetrameric state (8.6 S; 213 kDa) under apo conditions, as previously described^8^, or in the presence of GMP (8.7 S; 209 kDa). On the contrary, MgGTP induced the formation of an octamer (15.4 S; 460 kDa) as does the addition of 1 mM MgATP (14.8S; 463 kDa). Interestingly, the sedimentation distribution peak of MgGTP-bound form of IMPDHbt is shifted to higher sedimentation values in comparison with MgATP condition, suggesting a slightly less elongated and more compact conformation in the presence of this allosteric effector.

**Figure 3.**
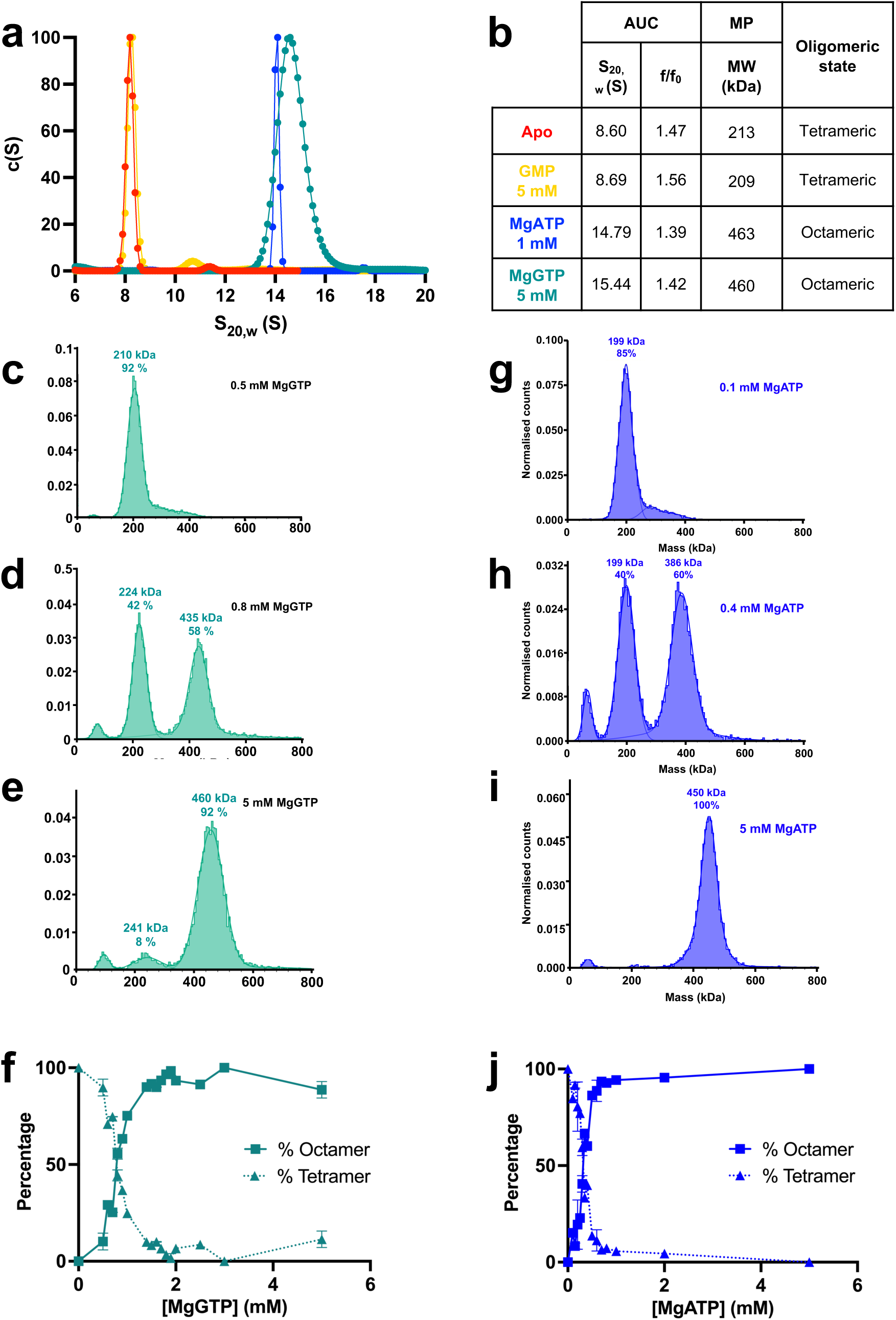
Regulation of the IMPDHbt quaternary structure by the substrates and the effectors. **a** Continuous c(S) distribution profiles as a function of the sedimentation coefficient normalized to standard conditions (S₂₀,w), measured under apo conditions (red) and in the presence of 5 mM GMP (yellow), 1 mM MgATP (blue), or 5 mM MgGTP (green). **b** Biophysical parameters of IMPDHbt determined by AUC (**a**) and MP (**c-j**) under apo conditions or in the presence of various effectors. S₂₀,w: sedimentation coefficient; f/f₀: frictional ratio; MW: molecular weight. **c-e** Normalized hit distribution as a function of molecular weight for IMPDHbt in the presence of increasing MgGTP concentrations (**c**, 0.5 mM; **d**, 0.8 mM; **e**, 5 mM). The percentage and molecular weight of each oligomeric species are indicated above each peak. **f** Quantification of the tetrameric (dotted line) and octameric (solid line) species as a function of MgGTP concentration. Percentages were calculated based on the number of hits corresponding to each oligomeric state. **g-i** Same as in **c-e** in the presence of increasing concentration of MgATP (**g**, 0.1 mM; **h**, 0.4 mM; **i**, 5 mM). **j** Same as in **f** as a function of MgATP concentration.

To better understand the relationship between the kinetic effects observed in the presence of MgATP and MgGTP, we monitored the change in the oligomeric state of IMPDHbt as a function of MgATP or MgGTP (Fig. 3c-j) concentrations using MP. These experiments revealed a concentration-dependent transition from tetramers to octamers upon the addition of either nucleotide. The estimated effective concentration required for 50 % octamerization (EC₅₀) was 0.8 mM for MgGTP (Fig. 3f) and 0.35 mM for MgATP (Fig. 3j). Thus, both nucleotides promoted the tetramer-to-octamer transition, but MgATP interacts with higher apparent affinity and more efficiently triggers the conformational shift.

Small angle x-ray scattering (SAXS) experiments (Fig. 4 and Supplementary Table 2) were performed to further confirm the oligomeric states and the distinct shapes observed by AUC and MP and to characterize the overall 3D-structure of IMPDHbt in solution with or without different ligands, in particular MgGTP and MgATP. The intensity curves, after background subtraction, displayed distinct variations depending on the experimental conditions (Fig. 4a). Guinier analysis confirmed monodisperse, non-aggregated samples (Fig. 4b), with radii of gyration of 44-53 Å, indicating a compact conformation. Normalized Kratky plots show a bell-shaped peak, consistent with a globular, folded enzyme, with slight shifts in apo, IMP, and GMP conditions suggesting increased quaternary flexibility. A shoulder characteristic of multi-domain proteins is observed in all conditions, accentuated by MgGTP and MgATP although differently, implying possible octamer conformational changes. Pair-distance distribution functions P(r) reveal a compact, globular structure (D_max_ 146-152 Å), although apo, IMP, and GMP states show broader distributions indicative of a more elongated conformation, consistent with R_g_ values from Guinier analysis (Fig. 4d). Collectively, AUC, MP, and SAXS analyses consistently revealed that both MgATP and MgGTP promote the tetramer to octamer transition of IMPDHbt, confirming the robustness of our observations across multiple structural approaches.

**Figure 4.**
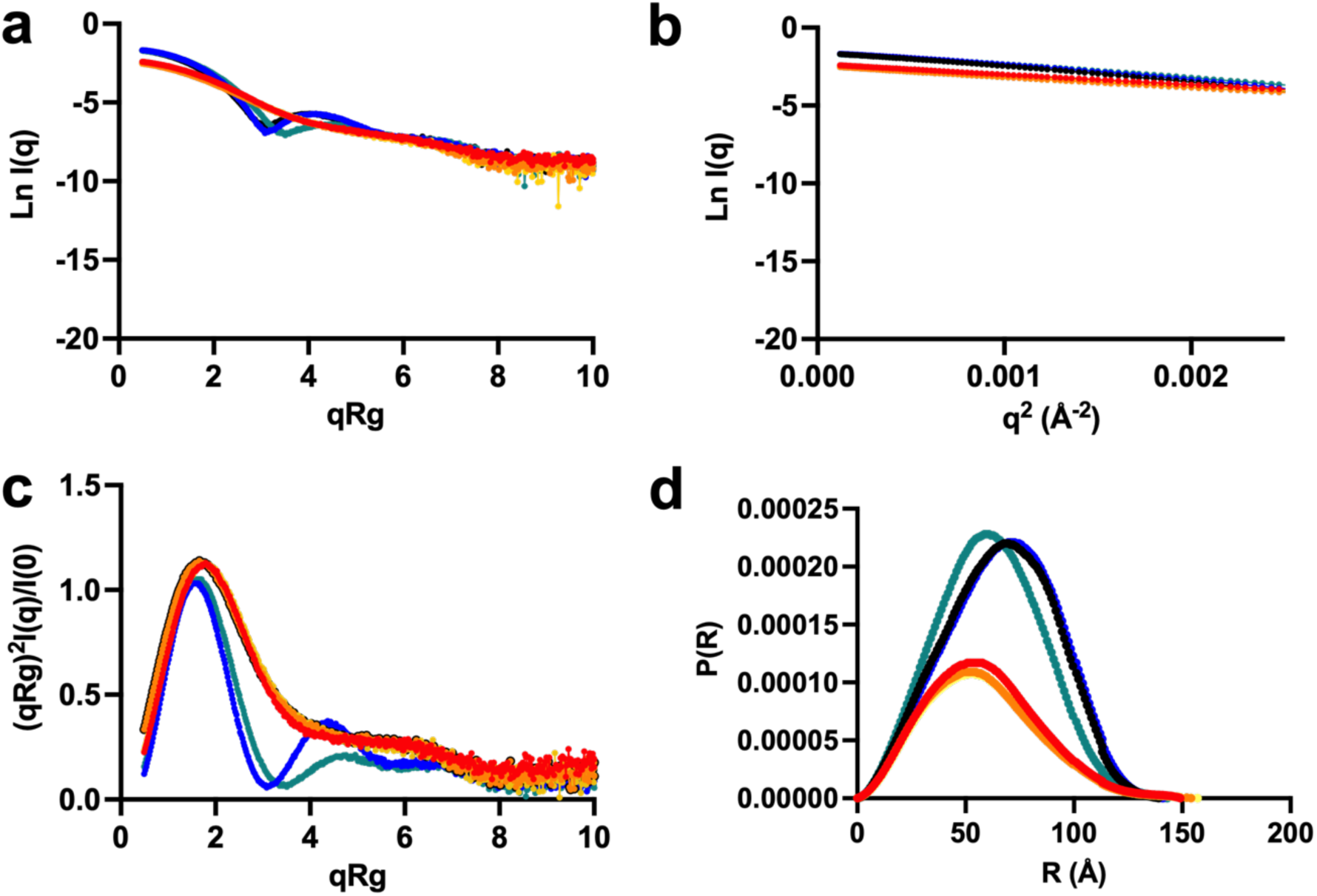
SAXS analysis of IMPDHbt conformational changes induced by different ligands. **a** scattering intensity curves, **b** Guinier plots, **c** normalized Kratky plots and **d** pair-distance distribution functions P(R)/I(0). Measurements were performed at a protein concentration of 3 mg/mL. Color code: apo (red), 6 mM IMP (orange), 5 mM GMP (yellow), 4 mM NAD^+^ (black), 5 mM MgATP (blue), and 5 mM MgGTP (green).

### Structural basis for ligand-dependent oligomerization of IMPDHbt

To elucidate the molecular mechanisms by which ligands and the regulatory Bateman domain control IMPDHbt activity and oligomerization, we determined several x-ray crystal structures that capture the enzyme in distinct, ligand-bound, and oligomeric states (see Table 2). These structures were solved to decipher (i) how ligand binding shape the active site and associated flexible loops, and (ii) how the Bateman domain mediates effector-dependent transitions between tetrameric and octameric assemblies. Across all structures, IMPDHbt adopted the canonical architecture of IMPDH enzymes, consisting of a catalytic (β/α)₈ barrel domain (or core domain) and a regulatory Bateman domain (Fig. 5a).

**Figure 5.**
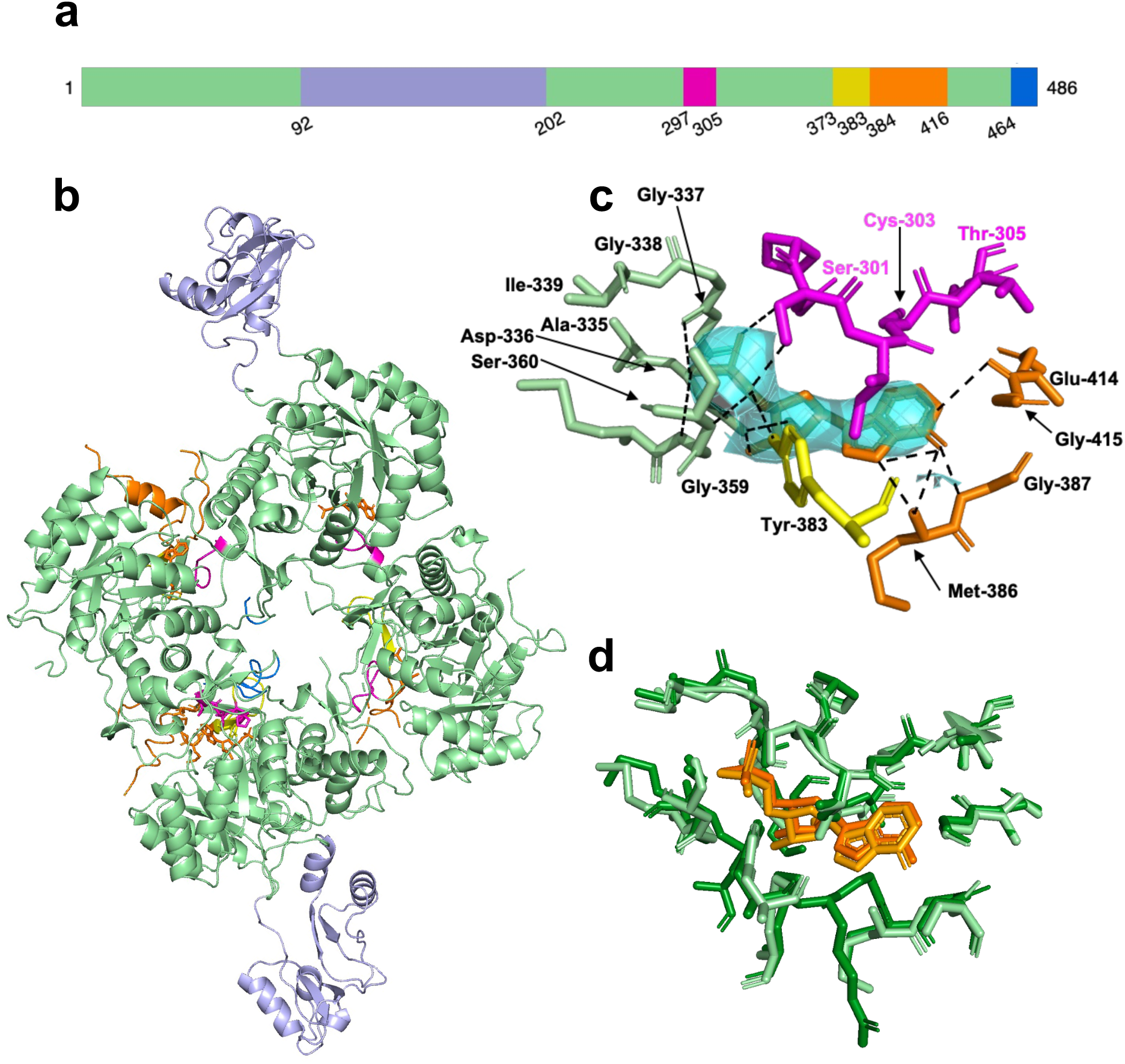
X-ray structure of IMPDHbt in the presence of IMP (PDB code 9HK6). **a** IMPDHbt primary sequence as a schematic bar representation. The core domain and the Bateman domain are coloured in light green and lavender, respectively, as well as important loops of the core domain involved in the enzymatic activity: catalytic loop (residues 297-305, pink), finger loop (residues 373-383, yellow), flap loop (residues 384-416, orange), and C-terminal loop (residues 464-486, blue) **b** Ribbon representation of the IMPDHbt tetramer bound to IMP (orange spheres), highlighting the Bateman domain, the core domain, and key structural loops: same colour code as in **a**. **c** Close-up view of the IMP-binding site. IMP is shown in orange with key interacting side chains in light green as ball-and-stick models. Hydrogen bonds are represented by black dashed lines. **d** Structural alignment of the IMP-binding site between IMPDHbt and the ΔBD variant of IMPDHpa (PDB code 5AHM). Side chains and IMP molecules are shown in ball-and-stick. For IMPDHbt, side chains and IMP are coloured as in **b**, while for IMPDHpa ΔBD the colours are shown in darker tones of green and orange, respectively.

**Table 2.** X-ray diffraction data collection and refinement statistics. IMPDHbt full-length in the presence of IMP (PDB code 9HK6), MgATP and MgGTP (PDB code 9HK7), MgGTP, MgATP, IMP and NAD^+^ (PDB code 9HK8) or MgATP, MgGTP and IMP (PDB 9HK9). IMPDHbt ΔBD variant in the apo form (PDB code 9HKA) or in the presence of GMP (PDB code 9HKB). Statistics for the highest-resolution shell are shown in parentheses.

| PDB id | 9HK6 | 9HK7 | 9HK8 | 9HK9 | 9HKA | 9HKB |
| --- | --- | --- | --- | --- | --- | --- |
| Wavelength | 0.9786 | 0.9786 | 0.9786 | 0.9793 | 0.9786 | 0.8266 |
| Resolution range | 49.3 - 3.35<br>(3.47 - 3.35) | 44.44 - 1.781<br>(1.845 - 1.781) | 37.71 - 2.75 (2.849 - 2.75) | 35.34 - 1.862<br>(1.928 - 1.862) | 41.94 - 2.154<br>(2.231 - 2.154) | 45.76 - 1.442<br>(1.494 - 1.442) |
| Space group | P 21 21 2 | I 4 | I 4 | I 4 | C 1 2 1 | P 4 |
| Unit cell | 137.58<br>197.2 93.19<br>90 90 90 | 119.6 119.6<br>143.61 90<br>90 90 | 119.24<br>119.24<br>141.98 90<br>90 90 | 118.96<br>118.96<br>141.84 90<br>90 90 | 136.7 72.92<br>95.71 90<br>125.59 90 | 91.51 91.51<br>86.32 90 90<br>90 |
| Total reflections | 507571<br>(80699) | 1302863<br>(136969) | 358350<br>(34395) | 1156908<br>(105649) | 293128<br>(28236) | 1773187<br>(176568) |
| **Unique reflections** | 37142<br>(3584) | 95888<br>(9544) | 25779<br>(2572) | 82113<br>(8091) | 40842<br>(3992) | 127652<br>(12631) |
| **Multiplicity** | 13.7 | 13.6 (14.4) | 13.9 (13.4) | 14.1 (13.1) | 7.2 (7.1) | 13.9 (13.9) |
| **Completeness (%)** | 99.74<br>(98.22) | 99.78<br>(99.95) | 99.95<br>(100.00) | 99.84<br>(98.63) | 98.32<br>(96.99) | 99.70<br>(99.27) |
| **Mean I/sigma (I)** | 13.41<br>(2.02) | 10.57<br>(2.90) | 13.14<br>(2.25) | 18.00<br>(1.86) | 16.93<br>(1.34) | 16.00<br>(1.63) |
| **Wilson B-factor** | 119.21 | 26.80 | 66.78 | 32.20 | 53.82 | 18.04 |
| **R-merge** | 0.156<br>(1.435) | 0.156<br>(1.034) | 0.1526<br>(1.347) | 0.1003<br>(1.283) | 0.05818<br>(1.344) | 0.07445<br>(1.259) |
| **R-meas** | 0.162<br>(1.488) | 0.1623<br>(1.072) | 0.1586<br>(1.401) | 0.1041<br>(1.336) | 0.06283<br>(1.451) | 0.0773<br>(1.307) |
| **R-pim** |  | 0.04442<br>(0.2824) | 0.04266<br>(0.3828) | 0.02761<br>(0.3678) | 0.02348<br>(0.5423) | 0.02064<br>(0.3494) |
| **CC1/2** | 99.9 (70.7) | 0.996<br>(0.766) | 0.998<br>(0.669) | 0.999<br>(0.743) | 0.999<br>(0.715) | 1 (0.866) |
| **CC\*** |  | 0.999<br>(0.931) | 0.999<br>(0.896) | 1 (0.924) | 1 (0.913) | 1 (0.963) |
| **Reflections used in refinement** | 37127<br>(3584) | 95874<br>(9539) | 25778<br>(2572) | 82103<br>(8087) | 40816<br>(3992) | 127362<br>(12630) |
| **Reflections used for R-free** | 1857 (179) | 4710 (466) | 1233 (100) | 4105 (404) | 2041 (200) | 6434 (633) |
| **R-work** | 0.2128<br>(0.2703) | 0.2888<br>(0.3506) | 0.1874<br>(0.3246) | 0.1761<br>(0.3534) | 0.1970<br>(0.3289) | 0.1608<br>(0.2954) |
| **R-free** | 0.2633<br>(0.3265) | 0.2956<br>(0.3499) | 0.2163<br>(0.3542) | 0.2141<br>(0.3947) | 0.2185<br>(0.3486) | 0.1744<br>(0.2963) |
| **CC(work)** | 0.866<br>(0.731) | 0.885<br>(0.583) | 0.967<br>(0.769) | 0.976<br>(0.821) | 0.959<br>(0.821) | 0.973<br>(0.878) |
| **CC(free)** | 0.787<br>(0.682) | 0.900<br>(0.563) | 0.918<br>(0.754) | 0.951<br>(0.765) | 0.950<br>(0.738) | 0.970<br>(0.850) |
| **Number of non-hydrogen atoms** | 11415 | 6930 | 7162 | 7831 | 4716 | 5852 |
| **macromolecules** | 11323 | 6291 | 6857 | 6834 | 4626 | 5240 |
| **ligands** | 92 | 130 | 266 | 201 | 0 | 76 |
| **solvent** | 0 | 509 | 39 | 796 | 90 | 560 |
| **Protein residues** | 1549 | 843 | 923 | 901 | 631 | 711 |
| **Nucleic acid bases** |  |  |  |  |  |  |
| **RMS(bonds)** | 0.002 | 0.003 | 0.003 | 0.006 | 0.002 | 0.014 |
| **RMS(angles)** | 0.57 | 0.61 | 0.61 | 1.02 | 0.43 | 1.32 |
| **Ramachandran favored (%)** | 94.81 | 96.03 | 96.61 | 97.65 | 96.60 | 97.69 |
| **Ramachandran allowed (%)** | 5.06 | 3.85 | 3.39 | 2.35 | 2.92 | 2.31 |
| **Ramachandran outliers (%)** | 0.13 | 0.12 | 0.00 | 0.00 | 0.49 | 0.00 |
| **Rotamer outliers (%)** | 0.00 | 0.45 | 4.71 | 0.82 | 1.05 | 1.48 |
| **Clashscore** | 8.23 | 4.94 | 4.41 | 4.16 | 2.25 | 3.48 |
| **Average B-factor** | 127.35 | 29.50 | 68.10 | 34.74 | 73.28 | 23.64 |
| **macromolecules** | 127.36 | 29.24 | 68.00 | 34.19 | 73.56 | 22.32 |
| **ligands** | 125.52 | 28.27 | 72.53 | 31.39 |  | 18.06 |
| **solvent** |  | 32.94 | 56.31 | 40.30 | 59.08 | 36.57 |
| **Number of TLS groups** | 20 | 12 | 8 | 9 | 8 |  |

### Tetrameric states reveal conserved active-site architecture and intrinsic regulatory flexibility

A tetrameric structure (PDB code 9HK6) captures full-length IMPDHbt in the presence of its substrate IMP in a catalytically primed configuration. IMP is positioned in the active site of each monomer, with the Cys303 oriented toward the C2 carbon of IMP and poised to form a covalent bond (Fig. 5). The substrate engages in conserved interactions involving residues critical for substrate recognition and catalysis. The atoms N7, O6, and N1 of the inosine nucleobase are stabilized by interactions with the backbone atoms of Met386, Gly387, and Glu414, respectively - all residues located in the “finger” loop. Tyr383, which is conserved across IMPDH species, mainly interacts with Gly359 and Ser360, while also forming, like Ser360, hydrogen bonds with the phosphate group of IMP. These interactions stabilize IMP within the active site and are accompanied by local rearrangements of loops, particularly the catalytic loop (Fig.5c, in pink), which adopts a closed conformation to accommodate the bound IMP molecule.

The orientation and binding mode of IMP in the active site (Fig. 5d) corresponds to that previously described in other IMPDHs, notably in the structure of IMPDHpa ΔDB (PDB code 5AHM^11^) or *Mycobacterium smegmatis* IMPDH (PDB code 8QQV^12^) in the presence of IMP. These results show that the IMP-binding site is well conserved among both IMPDH classes. Beyond the catalytic domain, this tetrameric structure also provided the first visualization of the Bateman domain in a non-octameric context. Remarkably, analysis of the tetrameric form revealed partial disorder within the Bateman and C-terminal domains across several monomers, suggesting substantial intrinsic flexibility of these regions in the absence of octamer-stabilizing ligands. This conformational plasticity likely permits the Bateman domain and the flexible loops to undergo effector-driven conformational rearrangements and promotes the transition from tetrameric to octameric assemblies upon binding of MgATP or MgGTP.

Consistent with this interpretation, and to directly assess the contribution of the Bateman domain to the structural regulation of IMPDHbt, we determined high-resolution crystal structures of Bateman domain-deleted variant (ΔDB) in the absence of any ligand (apo condition, PDB code 9HKA) and in the presence of GMP (PDB code 9HKB). In contrast to the full-length IMPDHbt, which can adopt either tetrameric or octameric assemblies depending on ligand conditions, all ΔBD variants crystallized exclusively as tetramers under all tested conditions, irrespective of ligand binding (Supplementary Fig. 3). This observation underscores the pivotal role of the Bateman domain in mediating inter-tetramer interactions required for octamer assembly. This is also in agreement with our previous studies using AUC^3^ on three ΔDB variants belonging to the two IMPDH classes. Structural alignment with the full-length tetrameric IMP-bound form revealed a high degree of similarity in the core catalytic domain, indicating that removal of the Bateman domain does not perturb the intrinsic fold of the catalytic monomer or its tetrameric assembly (Supplementary Fig. 3d).

Despite this overall structural conservation, the structures of the ΔBD variant revealed marked differences in the ordering of flexible loop regions compared to their full-length counterparts (Supplementary Fig. 3). In the apo ΔBD structure (PDB code 9HKA), solved at 2.15 Å, the catalytic loop (residues 189–196) was fully resolved in the electron density map and adopted an open conformation (Supplementary Fig. 3a). The finger loop (264–274) is also visible but only partially ordered. On the other hand, the flap (residues 275–307) and the C-terminal (residues 360–377) loops remained unresolved (Supplementary Fig. 3a). This behaviour is consistent with previous observations in full-length apo IMPDHpa (PDB code 6GJV), suggesting that these regions are intrinsically flexible and become stabilized only upon ligand binding (PDB code 4DQW). In the GMP-bound ΔBD structure (1.45 Å resolution, PDB code 9HKB), GMP occupied the active site of each monomer in a binding mode highly similar to IMP. The binding of GMP induced the closure of the well-defined catalytic loop in all monomers, consistent with productive substrate binding and catalytic readiness (Supplementary Fig. 3b). Notably, the flap loop remained largely unresolved in all ΔBD structures, suggesting that its stabilization is critically dependent on the presence of the Bateman domain, possibly via long-range allosteric interactions or direct interdomain contacts. Conserved interactions were identified between the guanine base and residues from the “finger” loop (Met277, Gly278, and Glu305), while the phosphate group engaged Tyr274 and Ser251 (Supplementary Fig. 3c). Although the residue numbers differ, these positions correspond structurally to the same elements that engage IMP in the full-length enzyme, with the renumbering resulting from deletion of the Bateman domain from the primary sequence. A unique hydrogen bond between the amino group of GMP and Thr196, absent in the IMP bound IMPDHbt structure, was also observed. These structural features correlate with enzymatic data, confirming the competitive inhibitory role of GMP towards IMP.

Comparative analysis with both the apo and ligand bound ΔDB structure with the IMP-bound full-length enzyme underscores the critical role of ligand binding in promoting an active-site architecture compatible with catalysis (Supplementary Fig. 3d). Specifically, both GMP and IMP not only induce a closed conformation of the catalytic loop but also draw the finger loop into closer proximity to the substrate within the active site. In contrast, in the apo form, the “finger” loop remains displaced, likely due to the lack of stabilizing interactions with a bound ligand. Together, these tetrameric structures suggest that ligand binding locally organizes the catalytic machinery, triggering a coordinated rearrangement of flexible loop regions and ensuring precise positioning of active-site elements without requiring large-scale domain movements. Overall, the tetramer emerges as a functional but highly flexible state, providing the conformational plasticity necessary for effector-driven octamer assembly.

### Effector-Bound octamers highlight flexible loop dynamics and combined ligand binding enhances structural rigidity

Crystallographic analyses in the presence of both MgATP and MgGTP revealed similar but distinct octameric assemblies of full-length IMPDHbt, each composed of two associated tetrameric rings, regardless of IMP presence (PDB code 9HK9, + IMP; PDB code 9HK7, - IMP). These two octamers differed in the organization of three flexible loops: the catalytic, finger and flap loops (Fig. 5a and 6). They exhibit varying degrees of ordering across the different conditions. Importantly, both the finger and flap loops contribute significantly to the interface between the two tetrameric rings that assemble into the octameric form. Their conformational status thus appears to correlate with the stability of the octamer, suggesting that these regions not only respond to effector binding but also act as structural determinants of oligomeric organization.

**Figure 6.**
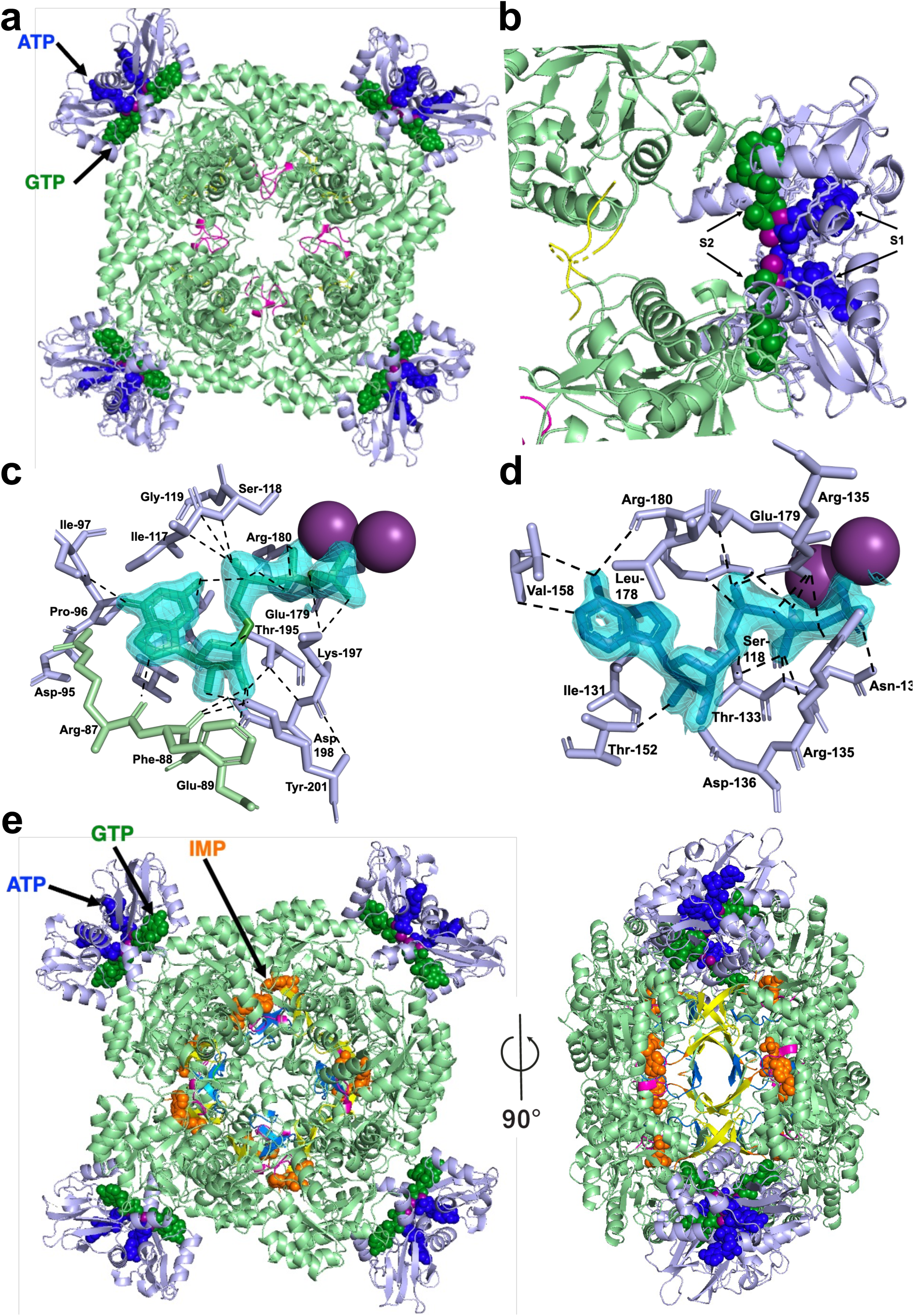
X-ray structures of IMPDHbt in the presence of MgATP and MgGTP without (PDB code 9HK7) or with IMP (PDB code 9HK9). **a** Ribbon representation of the IMPDHbt octamer bound to MgATP (blue spheres) and MgGTP (green spheres). The Bateman domain, the core domain, and key structural loops are coloured as depicted in Fig. 5a. The black arrow indicates one Bateman domain occupied by both MgATP and MgGTP. **b** Ribbon view of a pair of Bateman domains showing the S1 (MgATP-bound) and S2 (MgGTP-bound) sites. **c-d** Close-up views of the MgGTP- and MgATP-binding pockets in S1 and S2 sites, respectively. Interacting residues are shown as ball-and-stick models in light green (catalytic domain) and light blue (Bateman domain). Hydrogen bonds are depicted as black dashed lines. Mg²⁺ ions are shown as purple spheres. **e** Ribbon representation of the IMPDHbt octamer viewed along the 4-fold axis (left panel) or perpendicular to it (right panel) in complex with ATP (blue), GTP (green), IMP (orange) and Mg²⁺ ions (violet). Black arrows indicate a representative monomer simultaneously occupied by IMP in the core domain, and MgATP and MgGTP in the Bateman domain.

The 1,8 Å resolution structure (PDB code 9HK7) in the presence of MgATP and MgGTP revealed substantial stabilization of the Bateman domain. Unlike IMP-bound tetramer form, where only half of the four Bateman domains were resolved, all Bateman domains adopt a well-defined position relative to the catalytic core in the presence of MgATP and MgGTP (Fig. 6a). In these octameric assemblies, ATP and GTP bind specifically to the S1 and S2 sites of the Bateman domain, respectively. Each nucleotide via their phosphate groups chelates one magnesium ion (Fig. 6b). In the S2 site (Fig. 6c), the guanine base of GTP adopts a *syn* conformation and forms hydrogen bonds through its N2, O6, and C8 atoms with Arg180, Leu193, Ile97, and Arg87. The ribose hydroxyls are stabilized by Lys201 and Asp198, while the phosphate groups are engaged in electrostatic and hydrogen-bond interactions with Arg180, Lys197, and a coordinating magnesium ion. Similarly, in the S1 site (Fig. 6d), the adenine base of ATP binds in an *anti*-conformation, with the N6 atom forming hydrogen bonds with Val158 and Arg156. Ribose hydroxyl groups are recognized by the conserved Asp136 and Thr152, and the phosphate moieties interact with Glu179, Arg135, Arg180, and a magnesium ion. Similar ATP-GTP bound complexes have been reported in other IMPDHs (PDB codes: 8QQP^12^, 8G9B^13^, 6UA4^14^, 6UC2^14^, 6UA2^14^ and 7RGD^15^), though at lower resolution. These structures likewise show that the Bateman domain can bind ATP and GTP simultaneously and that the S2 site can accommodate either ATP or GTP, consistent with the binding mode observed in our high-resolution octameric assemblies.

Concerning the core domain, the catalytic loop is fully resolved and adopts a closed conformation, whereas other flexible elements (flap and C-terminal loops), apart from a portion of the finger loop, remain largely disordered. Interestingly, the octamer adopted a conformation, markedly different from that observed in the IMPDHpa in complex with MnATP complex (PDB code 4DQW) (Supplementary Fig. 4). This compactness might contribute to allosteric inhibition by restricting loop motions necessary for catalysis.

The presence of IMP in conjunction with ATP and GTP yielded an octameric assembly characterized by a more ordered finger loop at the intra-tetramer interface, while other elements, such as the flap loop, remain partially disordered (Fig. 6e). These findings suggest that IMP binding improves organization of the catalytic site but does not fully overcome the conformational constraints imposed by nucleotide-induced octamerization.

### A fully loaded active site captured in the presence of NAD⁺

Despite extensive efforts, obtaining a high-resolution IMPDHbt structure with clear electron density for a bound NAD⁺ molecule has proven challenging. Remarkably, only one condition led to successful visualization of NAD⁺: co-crystallization of IMPDHbt in the presence of IMP, NAD⁺, MgGTP, and MgATP. This octameric structure (PDB code 9HK8), resolved at 2.75 Å, reveals a fully loaded active site with one molecule of NAD⁺ and IMP per monomer, alongside two magnesium ions, one ATP and one GTP molecules bound to the Bateman domain (Fig. 7). The NAD⁺ molecule adopted a conserved binding mode analogous to that described for *T. foetus* IMPDH^16^. Key hydrogen bonds are observed between the hydroxyl groups of the nicotinamide ribose and Asp246, as well as between the adenosine ribose and Ser252. Additional stabilization arises from interactions between the NAD⁺ carboxamide group and the peptide backbone of Gly296, Gly298, and Ile297. The adenosine moiety was nestled in a hydrophobic pocket between Pro24 and His249, ensuring tight anchoring of the cofactor. The nicotinamide ring of NAD⁺ aligned precisely with the base moiety of IMP, ensuring an optimal arrangement that positions the hydride donor and acceptor in an ideal geometry for catalysis (Fig. 7c). Also, in this structure, most of the previously unresolved regions became visible. Residues 383–395 of the flap loop, absent in all previous structures, are now resolved, although the loop spanning residues 394–411 remains partially disordered. In addition, the C-terminal loop (residues 464–480) is nearly fully resolved, with only the last seven residues missing. The simultaneous presence of IMP, NAD⁺, ATP, and GTP appears to drive the observed rigidification effect. The combined binding appears to stabilize mobile regions by engaging the catalytic and regulatory domains synergistically, thereby locking the enzyme in a catalytically competent state. Together, this structural snapshot offers a rare view of the fully assembled catalytic site. The cooperative stabilization of mobile elements by multiple ligands suggests a finely tuned interplay between substrate binding and allosteric regulation, and provides molecular insights into the control of enzymatic activity in response to intracellular nucleotide pools.

**Figure 7.**
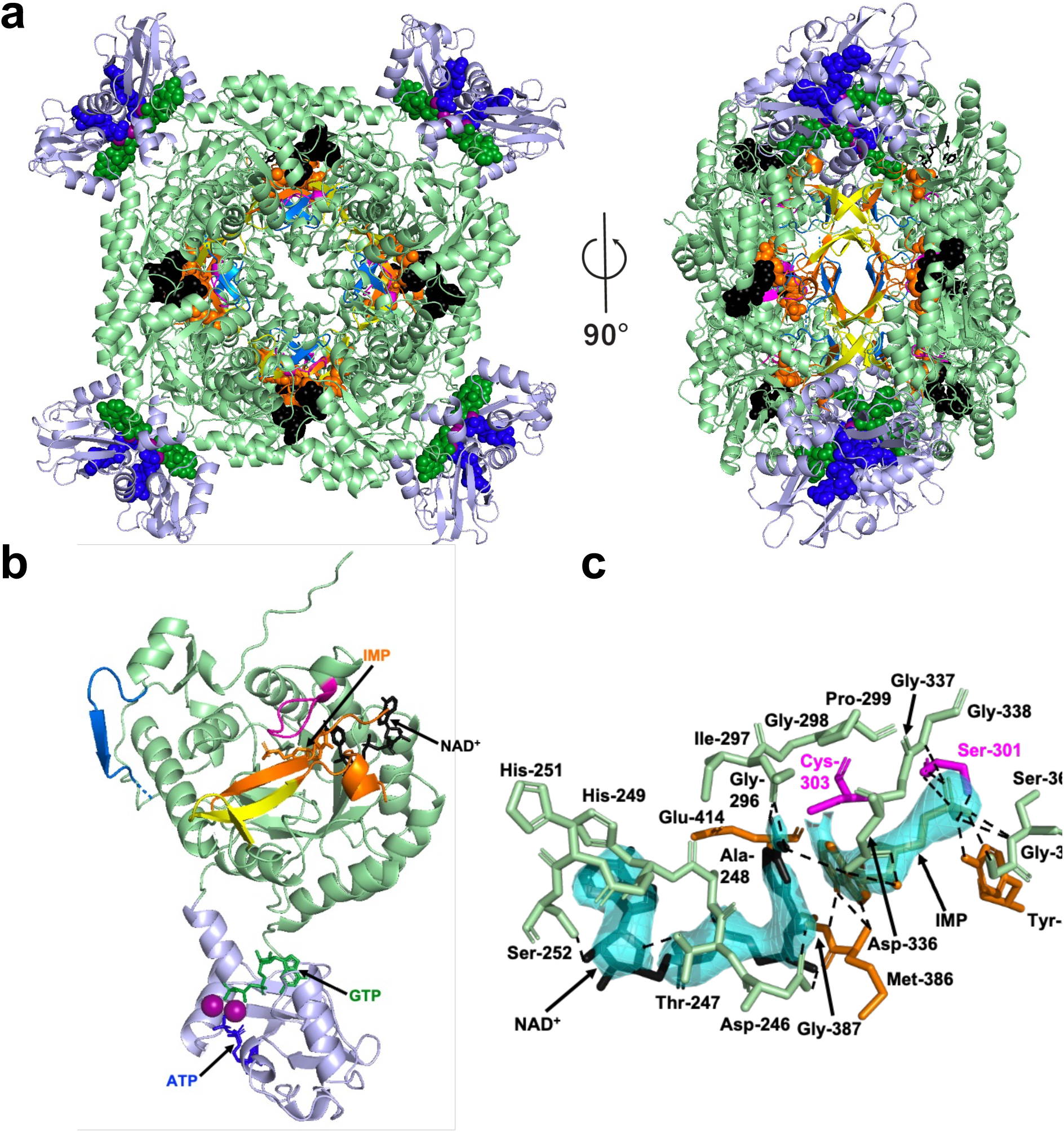
X-ray structures of IMPDHbt in the presence of IMP, NAD^+^, MgATP and MgGTP (PDB code 9HK8). **a** Ribbon representation of the octameric IMPDHbt viewed along the 4-fold axis (left panel) or perpendicular to it (right panel) in complex with IMP (orange), NAD⁺ (black), ATP (blue), GTP (green) and Mg²⁺ ions (violet). The Bateman domain, the core domain, and key structural loops are coloured as depicted in Fig. 5a. **b** Ribbon view of a single IMPDHbt monomer showing ligand binding: IMP and NAD⁺ in the active site, MgATP and MgGTP in the canonical S1 and S2 sites of the Bateman domain. **c** Close-up of the NAD⁺-binding site. NAD⁺ is shown in black and side chains of interacting residues are shown as ball-and-stick models in light green (catalytic domain), pink (catalytic loop) and orange (flap loop). IMP is shown in orange. Hydrogen bonds are indicated by black dashed lines.

**Figure 8.**
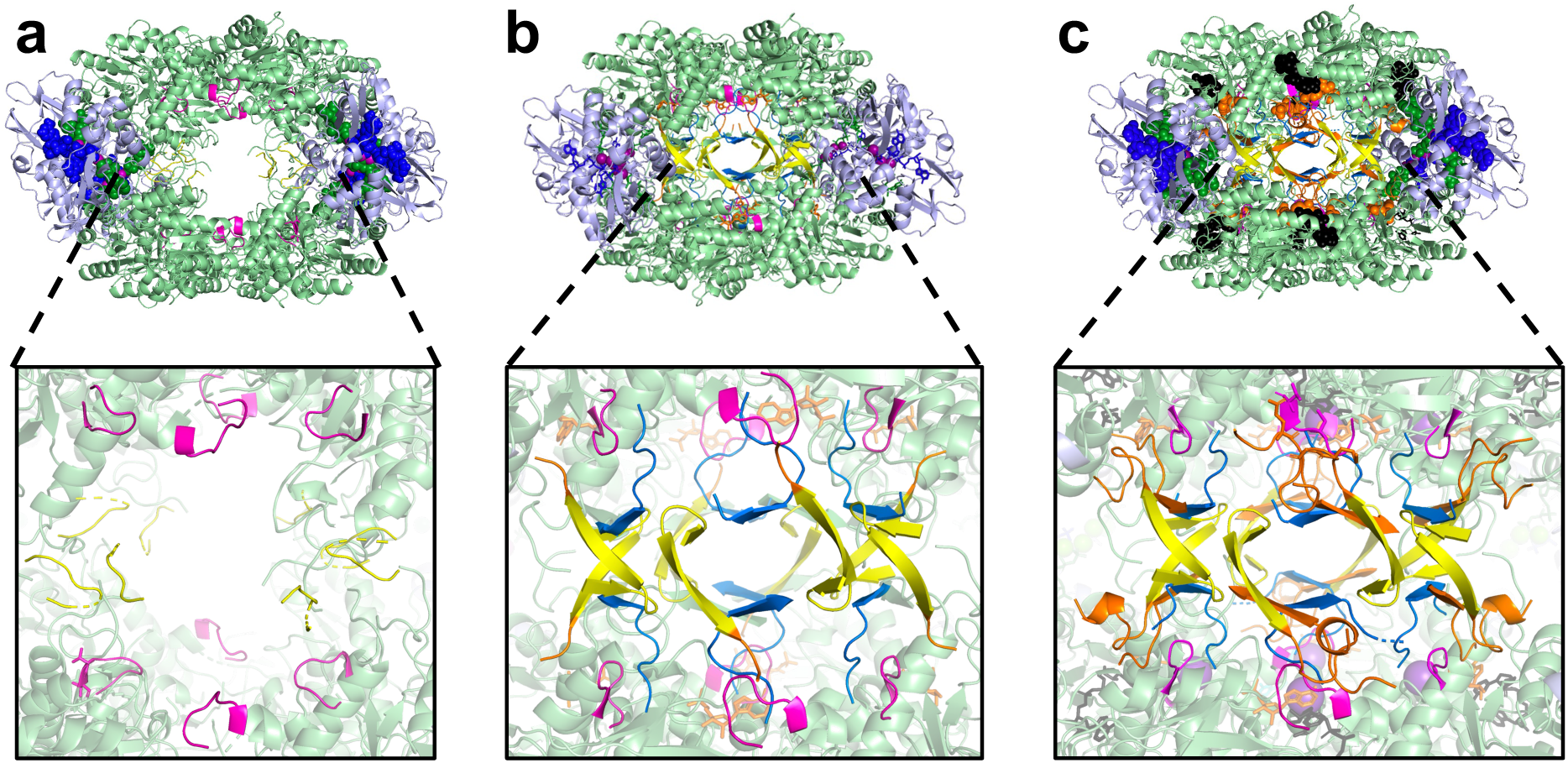
Structural reorganization of loops in the IMPDHbt core domain. Comparison of the three octameric 3D structures of IMPDHbt (**a** PDB code 9HK7, **b** PDH code 9HK8 and **c** PDB code 9HK9), with close-up views of important loops (same colour code as in Fig. 5a).

## Discussion

This study provides the first comprehensive characterization of IMPDHbt, a representative of class II bacterial IMPDHs and establishes a mechanistic framework for understanding how nucleotides regulate both the enzyme’s catalytic activity and oligomeric state. By combining enzymology and integrative structural biology, we demonstrate how ligand binding is coupled to functional and structural transitions within the enzyme.

A central finding of this work is the identification of MgGTP as a direct and independent allosteric inhibitor of IMPDHbt. In addition to GMP and XMP, which also inhibit catalytic activity, MgGTP acts as a potent negative effector even in the absence of MgATP, significantly reducing catalytic activity with an IC₅₀ in the millimolar range. This contrast with previous work by Fernandez-Justel *et al.*^17^ in which they reported that GTP-mediated allosteric inhibition requires the presence of ATP and instead reveals an intrinsic sensitivity of class II IMPDHs to guanine nucleotide levels. When both MgATP and MgGTP are present, the enzyme exhibits a biphasic response: sub-millimolar concentrations of MgGTP enhance activity in the presence of MgATP, whereas higher concentrations restore inhibition. This behavior highlights a sophisticated regulatory mechanism in which GTP inhibits the enzyme on its own, while ATP fine-tunes the response rather than simply counteracting it. Together, these nucleotides act as coordinated regulatory signals that enable the enzyme to sense and respond to fluctuations in the intracellular adenine-to-guanine nucleotide ratio. This coordinated modulation reveals an intricate interplay between positive and negative allosteric effectors, a relatively uncommon mechanism in enzyme regulation, where ligands typically compete for the same site rather than acting in a coordinated, concentration-dependent manner. In this context, IMPDH emerges as a metabolic regulator that dynamically adjusts guanine nucleotide biosynthesis in response to cellular demands.

To understand the structural basis of this regulation, our combined biophysical analysis using AUC, MP and SAXS demonstrate the crucial role of the Bateman domain in mediating regulation by ATP and GTP. Consistent with these observations, the ΔDB variant remains strictly tetrameric under all tested conditions and loses all responsiveness to MgGTP, while retaining sensitivity to GMP. This strongly supports the conclusion that the Bateman domain serves as the exclusive sensor of nucleotide triphosphate-mediated regulation of IMPDHs. Notably, this regulation is mediated by a single regulatory domain, which integrates opposing nucleotide signals to generate distinct functional outputs depending on ligand identity and concentration.

In the absence of effectors or in the presence of GMP or IMP, IMPDHbt adopts a tetrameric state. The binding of MgGTP or MgATP induces a transition toward an octameric assembly. Quantitative analysis further shows that MgATP induces this transition with higher apparent affinity (EC₅₀ ≈ 0.35 mM) compared to MgGTP (EC₅₀ ≈ 0.8 mM). Importantly, although one might expect GTP-mediated inhibition to shift the equilibrium back toward the tetrameric state, our results demonstrate that the enzyme remains predominantly octameric even at saturating MgGTP concentrations. Our SAXS and x-ray data further indicates that inhibition arises from subtle conformational transitions within the octameric assembly rather than from a change in oligomeric state itself. Specifically, although both MgATP and MgGTP promote tetramer-to-octamer conversion, inhibition correlates with changes in overall compaction and global shape of the protein as well as rigidification of crucial loops involved in the catalytic reaction, including the catalytic, finger, and flap loops, which are essential for substrate turnover. These observations support a model in which distinct octameric conformations encode different functional states, where ligand-specific rearrangements of flexible elements modulate active-site accessibility and dynamics. In this context, the Bateman domain emerges as a conformational switch that transduces nucleotide binding into long-range structural effects.

On the other hand, the physiological significance of the tetrameric state (observed in the apo form or when bound to GMP/IMP) remains unclear. Particularly, given that intracellular ATP and GTP concentrations are typically in the millimolar range^18^, the octameric assembly likely represents the predominant physiological species *in vivo*. However, the conservation of the tetrameric form for class II IMPDHs raises the possibility that this oligomeric state may fulfill additional roles beyond canonical IMPDH catalytic activity. Indeed, several IMPDHs from different organisms have been reported to exhibit moonlighting functions, ranging from transcriptional regulation (*Drosophila melanogaster* IMPDH^19^), to nucleic acid binding (human, *Tritrichomonas foetus* and *E. coli* IMPDHs^20, 21^). It is therefore tempting to speculate that the tetrameric state represents a functionally distinct structural platform, potentially dedicated to regulatory or scaffolding roles independent of catalysis.

In higher eukaryotes, this oligomeric versatility extends to the formation of higher-order assemblies^22, 23^: human IMPDH1^9, 15, 24, 25^ and IMPDH2^14, 26, 27^ can polymerize into extensive filamentous structures (historically termed “rods and rings”, now designated IMPDH filaments). These structures assemble dynamically in response to metabolic stress (such as glutamine deprivation, serine limitation, or inhibition of GMP synthase) and disassemble upon restoration of guanine nucleotide levels. Recent work has shown that these filaments represent catalytically active polymers that resist allosteric feedback inhibition by GTP, thereby maintaining *de novo* GMP synthesis during periods of high purine demand in rapidly proliferating cells, including embryonic stem cells and activated T lymphocytes. In addition to their metabolic role, filament formation may also regulate the moonlighting functions of IMPDH. It has been proposed that polymerization could sequester the interaction of the enzyme with nucleic acids or modulate its nucleocytoplasmic shuttling, thereby controlling its non-catalytic activities such as DNA binding or transcriptional regulation. Moreover, these assemblies may perturb the formation of the supramolecular complex constituted by the enzymes involved in the *de novo* purine nucleotides biosynthesis, including IMPDH. This complex has been demonstrated to form a metabolon called purinosome in mammals enabling coordination of the enzymatic activities and enhancing metabolic flux under conditions of high nucleotide demand^28, 29^. In contrast to the filament-based regulation observed in human IMPDHs, our results highlight a distinct mechanism in bacterial IMPDHbt, where fine-tuned modulation arises from the interplay between ATP and GTP at the level of the Bateman domain, rather than from large-scale polymerization.

To gain molecular insight into the IMPDH regulatory mechanism, we solved six high-resolution crystal structures capturing distinct functional states of the full-length or ΔBD variant. Importantly, one structure provides a rare snapshot of a fully assembled pre-catalytic complex, in which IMP and NAD⁺ are simultaneously bound while ATP and GTP occupy their respective allosteric sites. Previous structural studies have described IMPDH in distinct catalytic or regulatory states, but never with catalytic substrates bound under allosteric control. For example, Johnson *et al.*^14^ visualized human IMPDH2 filaments stabilized by ATP/GTP (e.g., PDB 6U8E), revealing the architecture of the regulatory polymer but at resolutions (3.1 - 4.2 Å) insufficient to resolve substrates within the active site. Two other crystallographic structures (PDB codes 1MEW and 4ZQM from *Tritrichomonas foetus* IMPDH^16^ and *Mycobacterium tuberculosis* ΔDB IMPDH, respectively) depict product-bound states with XMP and NAD⁺, corresponding to the post-catalytic endpoint following hydride transfer and covalent intermediate hydrolysis. In contrast, our structure captures a rare pre-catalytic Michaelis complex [E·IMP·NAD⁺] that coexists with MgATP and MgGTP bound at their allosteric sites. This state arises from a crystallization condition containing Na⁺ (50 mM NaCl) but lacking K⁺, under which IMPDHbt is catalytically inert despite adopting a catalytically competent conformation. As previously reported, K⁺ is an essential cofactor for the enzyme to display maximal activity ⁺ (∼100 mM)^8^, reflecting an absolute requirement for this cation. In its absence, productive hydride transfer between IMP and NAD⁺ cannot proceed, as the K⁺-dependent activation of the water molecule required for hydrolysis of the covalent E–XMP* intermediate is impaired. As a result, the catalytic cycle is arrested prior to redox chemistry, allowing direct visualization of the enzyme poised for catalysis within an allosterically regulated context. This structure therefore provides a unique crystallographic snapshot of the catalytic machinery aligned for hydride transfer under physiologically relevant regulatory conditions.

Beyond these mechanistic insights, our findings have important implications for antimicrobial strategies. A variety of natural allosteric regulators have been characterized for several bacterial IMPDHs^1^, reflecting adaptation to diverse physiological conditions. These chemically diverse compounds bind to the Bateman domain through multiple binding pockets. Besides ATP and GTP, other nucleotides have been shown to inhibit IMPDHs, particularly in bacterial enzymes. Notably, the alarmone (p)ppGpp, which regulates the stringent response to nutritional stress, was identified as a direct allosteric inhibitor of IMPDH in species, such as *Bacillus subtilis* and *Streptomyces coelicolor*^17^. Likewise, diadenosine tetraphosphate (Ap4A), a putative second messenger, inhibits *B. subtilis* IMPDH upon binding to its Bateman domains^30^, but has no effect on the *E. coli* enzyme^31^. These variations in regulatory mechanisms likely reflect adaptations to different environments and stress conditions. The plasticity of the Bateman domain, together with its ability to accommodate diverse ligands, highlights this region as a promising target for the development of selective allosteric inhibitors for bacterial IMPDHs. Unlike active-site inhibitors, which often suffer from poor specificity, targeting the Bateman domain could enable species-specific modulation of IMPDH activity, particularly in pathogenic bacteria such as *Burkholderia*. In this context, four synthetic chemical series were identified as allosteric inhibitors of IMPDHpa through the screening of compound libraries^32^, highlighting its druggability. Co-crystallization experiments of IMPDHpa with one of these molecules (PDB code 6GK9) showed a structure identical to that in apo condition (PDB code 6GJV). Unlike MgATP, a single molecule of the allosteric inhibitor binds per Bateman domain at a site superimposed with the canonical S2 site, and this site is located on two adjacent monomers. The structural differences observed between ATP- and GTP-bound states further suggest that ligand-specific conformations could be exploited for rational drug design.

Finally, our results reinforce the view of IMPDH as a central regulator at the interface of metabolism and cellular signaling. Rather than operating through simple feedback mechanisms, IMPDH employs versatile and multilayered regulatory strategies than previously appreciated, integrating multiple nucleotide signals at both the catalytic and structural levels. This enables the enzyme to generate graded functional responses to changing cellular conditions. Our findings therefore extend the current paradigm of IMPDH regulation, positioning the enzyme not only as a key metabolic catalyst but also as a dynamic hub linking nucleotide homeostasis to broader physiological cues. More broadly, these results highlight the need to reassess IMPDH regulatory mechanisms across different organisms and physiological contexts.

## Methods

### Cloning and protein production and purification

Plasmids used in this work are listed in Supplementary Table 3. These constructs were obtained as described previously^3, 8, 9^. The experimental protocol employed for the expression and purification of the different recombinant protein closely followed the procedure previously outlined in Alexandre *et al.*^8^ and in Gedeon *et al.*^3^ (see the composition of the purification buffers in Supplementary Table 3). Briefly, the recombinant proteins (with a His-tag at the N-terminus) expressed in *E. coli* strain BL21(DE3)/pDIA17^33^ were purified using a two-step procedure involving affinity chromatography followed by size-exclusion chromatography.

### Enzymatic assay and kinetic characterization

The IMPDH catalytic activity was measured at 30°C by monitoring the NADH production at 340 nm on an Eppendorf ECOM 6122 photometer. All enzymatic reactions were carried out in a total volume of 0.5 mL of a corresponding optimal kinetics buffer noted buffer K (composition in Supplementary Table 3) previously determined by screening various buffer compositions adjusted to the appropriate pH and KCl concentrations. Initial velocity values were determined at various concentrations of IMP or NAD^+^, in the presence or absence of saturating concentrations of effectors (GMP, MgATP or MgCl_2_-GTP). Enzymatic reactions were initiated by adding IMPDH (final concentration: 0.1–2 µM), previously diluted in Buffer A (Supplementary Table 3) supplemented with 1 mM DTT and 1 mM EDTA. Saturating concentrations of IMP, NAD^+^, GMP, MgATP, and MgGTP were determined individually for each IMPDH variant. Experimental data were fitted using GraphPad Prism v10.3.1 (GraphPad Software, Inc.) and KaleidaGraph (Synergy Software, Inc.) according to one of the following equations, depending on the kinetic behaviour of the variant: i) Michaelis-Menten equation v = V_m_ [S]/(K_m_ + [S]); ii) the substrate inhibition equation v = V_m_ [S]/(K_m_ + [S] + [S]^2^/K_I_); iii) Hill equation v = V_m_ [S]^nH^ /(K_0.5_^nH^ + [S]^nH^), where v is the reaction rate, V_m_ the maximal rate, [S] the NAD^+^ or IMP concentration, K_m_ the Michaelis-Menten constant, K_I_ the inhibitory constant, K_0.5_ the IMP concentration at half-saturation, and n_H_ the Hill number index. IC_50_ values were determined according to the dose response logistic. One unit of enzyme activity corresponds to 1 µmole of the product formed in 1 min at 30°C under the assay conditions.

### Analytical ultracentrifugation (AUC)

Sedimentation velocity experiments were carried out at 20°C using a ProteomeLab XL-I analytical ultracentrifuge from Beckman-Coulter equipped with both UV-Vis absorbance and Rayleigh interference detectors. All experiments were performed on the Molecular Biophysics Core Facility at Institut Pasteur. Sample were prepared in buffer A (Supplementary Table 3) in which 0.5 mM tris(2-carboxyethyl)phosphine (TCEP) was used as a reducing agent in place of DTT. Experiments were performed under apo conditions (no ligand) or in the presence of saturating concentrations of ligands, as defined by prior enzymatic assays: 5 mM GMP, 1 mM MgATP, 5 mM MgGTP, and a mixture of 5 mM MgATP + 5 mM MgGTP. A consistent protein concentration of 1 mg/mL was chosen for all conditions. A total volume of 400 μL of samples was loaded into a two-sector epoxy centrepiece with sapphire windows, along with 420 μL of protein-free buffer. Samples were subjected to centrifugation at 54,000 rpm using an An-60 Ti rotor. The protein’s partial specific volume, and the viscosity and density of its corresponding buffer A (see Supplementary Table 3) were estimated using Sednterp 1.09, available online from The Boston Biomedical Research Institute. Raw data were analysed using Sedfit 12.0^34, 35^, employing a continuous size distribution c(s) model with interference optics to determine sedimentation coefficient distributions, normalized to standard conditions (20 °C, eau) and frictional ratios (f/f0).

### Mass Photometry

Molecular mass and oligomeric state analysis of IMPDHbt were conducted using the second generation of the Refeyn Two^MP^ instrument. The measurements adhered to the standard protocol described by Wu and Piszczek^36^, involving the dilution of protein preparations to the optimal concentration of 200 nM in the appropriate buffer A (Supplementary Table 3) supplemented with 0.5 mM TCEP, and filtered through 0.22 µm syringe filters prior to measurement. Measurements were conducted under various conditions: in the absence of ligands (apo) and in the presence of fixed concentrations of substrates and effectors, including 5 mM IMP, 5 mM NAD⁺, MgATP (0.1–5 mM), and MgGTP (0.5–5 mM). For each experiment, an 18 μL volume of ligand-specific buffer was applied to an isopropanol sonication-cleaned glass coverslip. The instrument automatically established focus, and just before data acquisition, 2 μl of the diluted sample were swiftly added into the sample well containing buffer and thoroughly mixed by pipetting, yielding a final protein concentration of 20 nM.

A one-minute recording of the collision events was performed using the AcquireMP software (Refeyn), and data were analyzed using the DiscoverMP software (Refeyn). The contrast of the landing events on the surface (point spread functions) was scrutinized and converted to molecular mass using two standard calibrants urease and BSA (Sigma-Aldrich) known for well-defined molecular mass peaks (91, 272, and 545 kDa and 66, 133, and 199 kDa, respectively) that facilitated the estimation of the molecular weight of particles within preparation. The number of molecular collisions was plotted against the measured molecular weight, and the peaks were fitted to a Gaussian distribution using DiscoverMP to estimate the oligomeric species present in the sample.

### Size exclusion chromatography small angle X-ray scattering

SAXS experiments were conducted at the SWING beamline at the Synchrotron SOLEIL in Saint-Aubin. A total volume of 50 µL of IMPDHbt protein at 3.14 mg/mL in buffer A supplemented with 1 mM DTT and 1 mM EDTA (50 mM Na_2_CO_3_ pH 9.5, 100 mM KCl, 1 mM DTT, 1 mM EDTA) was injected onto a size-exclusion chromatography column (Superose® 6 Increase 5/150 mm, Cytiva Life Sciences) pre-equilibrated at 4 °C. Experiments were performed in the absence or presence of saturating concentrations of ligands (6 mM IMP, 4 mM NAD⁺, 5 mM GMP, 5 mM MgATP, 5 mM MgGTP, or a mixture of 5 mM MgATP and 5 mM MgGTP). The column was directly coupled to the SAXS flow-through capillary cell for continuous sample delivery during data acquisition. Scattering data were recorded using the SWING beamline with a wavelength of 150–156 Å and processed using standard protocols. SAXS data were collected and the curves were background-subtracted using FOXTROT. Data were processed using standard procedures and were analyzed using the PRIMUS program^37^. Normalized Kratky plots were determined using the Guinier analysis, following the methodology outlined by Durand et al. (2010)^38^ and Pérez et al. (2001)^39^. The pair-distance distribution function P(r) were generated from the scattering curves using an indirect transform method in GNOM (Svergun, 1992)^40^. All these programs are found within the ATSAS software package^41^.

### Protein Crystallization and X-ray diffraction

High-throughput initial screening of crystallization conditions for IMPDHbt and IMPDHbt ΔDB was carried out using a MosquitoTM nanoliter-dispensing crystallization robot (TTP Labtech) and commercial crystallization screens (Hampton Research, Molecular Dimensions, Jena Biosciences, Emerald Biosystems). These experiments were performed using the protocols and the procedure established at the Crystallography Core Facility of the Institut Pasteur^42^. The best crystals were obtained in the crystallization conditions listed in Supplementary Table 4. Before cryocooling, the single crystals were soaked in a reservoir solution supplemented with a cryoprotectant listed in Supplementary Table 4.

X-ray diffraction data from flash-cooled single crystals were collected on the Proxima-1 or Proxima-2 beamlines at Synchrotron SOLEIL (Saint-Aubin, France). Structures were solved by molecular replacement using Phaser AutoMR with previously published IMPDHpa structures (PDB code 6GJV and 4DQW for the apo^32^ and MgATP-bound^9^ forms, respectively) as search models. Refinement was conducted with several types of refinement approaches using the current Python-based Hierarchical Environment for Integrated Crystallography (Phenix suite) software^43, 44, 45^ and with alternating manual rebuilding in COOT^46^. Final models all have favorable R / Rfree values, Molprobity scores and excellent geometry and stereochemistry. Data processing and model refinement statistics are shown in Table 2. Coordinates and structure factors have been deposited in the Protein Data Bank under accession codes 9HK6, 9HK7, 9HK8, 9HK9, 9HKA and 9HKB (Table 2). All structural figures were generated with the PyMol Molecular Graphics System, Version 1.3r1 (Schrödinger, LLC).

## Supporting information

Supplementary material - Ayoub et al.

## Acknowledgements

We thank the Institut Pasteur’s platforms for their support: the staff of the Crystallography platform, the “Plateforme de Milieu” for media culture preparation, and the Molecular Biophysics Platform for access to state-of-the-art instrumentation. We acknowledge SOLEIL (St. Aubin, France) for provision of synchrotron radiation facilities, and we would like to thank beamline staff for assistance in using beamlines PROXIMA-1, PROXIMA2A, and SWING. We thank Sarah Dubrac, Nicolas Wolff and Marc Feuilloley for fruitful discussions.

This work was supported in part by the Centre National de la Recherche Scientifique (CNRS), the Institut National de la Santé Et de la Recherche Médicale (INSERM) and the Institut Pasteur. Nour Ayoub acknowledges a PhD fellowship from Médicament, Toxicologie, Chimie et Imagerie Ph.D. school (MTCI, ED 563), Université Paris Cité.

## Author contributions

Nour Ayoub, and Hélène Munier-Lehmann conceptualized and designed experiments. Hélène Munier-Lehmann supervised the work. Nour Ayoub provided purified proteins, accomplished biochemistry experiments and biophysical characterization, and analyzed corresponding data. Nour Ayoub, and Bertrand Raynal performed SAXS experiments and conducted the SAXS data analysis. Nour Ayoub, and Ahmed Haouz conducted crystallization, and collected x-ray data; Muriel Gelin, and Gilles Labesse solved structures. Nour Ayoub, and Hélène Munier-Lehmann wrote the article with input from all authors.

