## Supplementary material - Ayoub et al. for "Deciphering the nucleotide driven kinetic and oligomeric dynamics in IMPDH regulation"

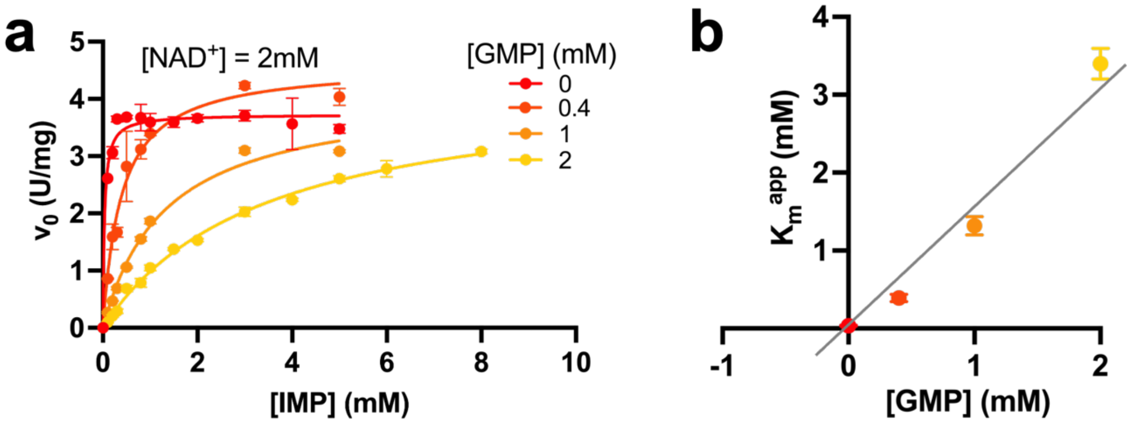

**Supplementary Fig. 1- IMPDHbt inhibition by GMP. a** Plots of the initial velocity of IMPDHbt as a function of IMP concentration in the presence of 2 mM $NAD^+$ and 0, 0.4, 1 or 2 mM GMP. Each data point represents a mean ± standard deviation in error bars. The curves correspond to the fit of the experimental data to the Michaelis–Menten equation. **b** Plot of the apparent $K_m^{IMP}$ deduced from the curves in panel **a** as a function of GMP concentration was used to determine the $K_i$ for GMP.

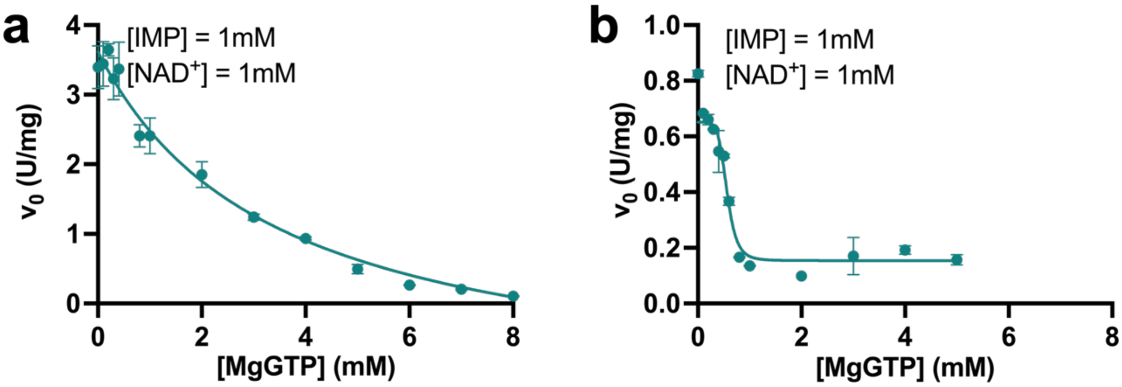

**Supplementary Fig. 2- IMPDHec and IMPDHpa inhibition by MgGTP. a** Plots of the initial velocity of IMPDHec (**a**) and IMPDHpa as a function of MgGTP concentration in the presence of 1 mM IMP and 1 mM $NAD^+$. The curves correspond to the fit of the experimental data to the following equation Y = Bottom + (Top-Bottom)/(1+(X/IC50)) and the calculated parameters are displayed in Supplementary Table 1.

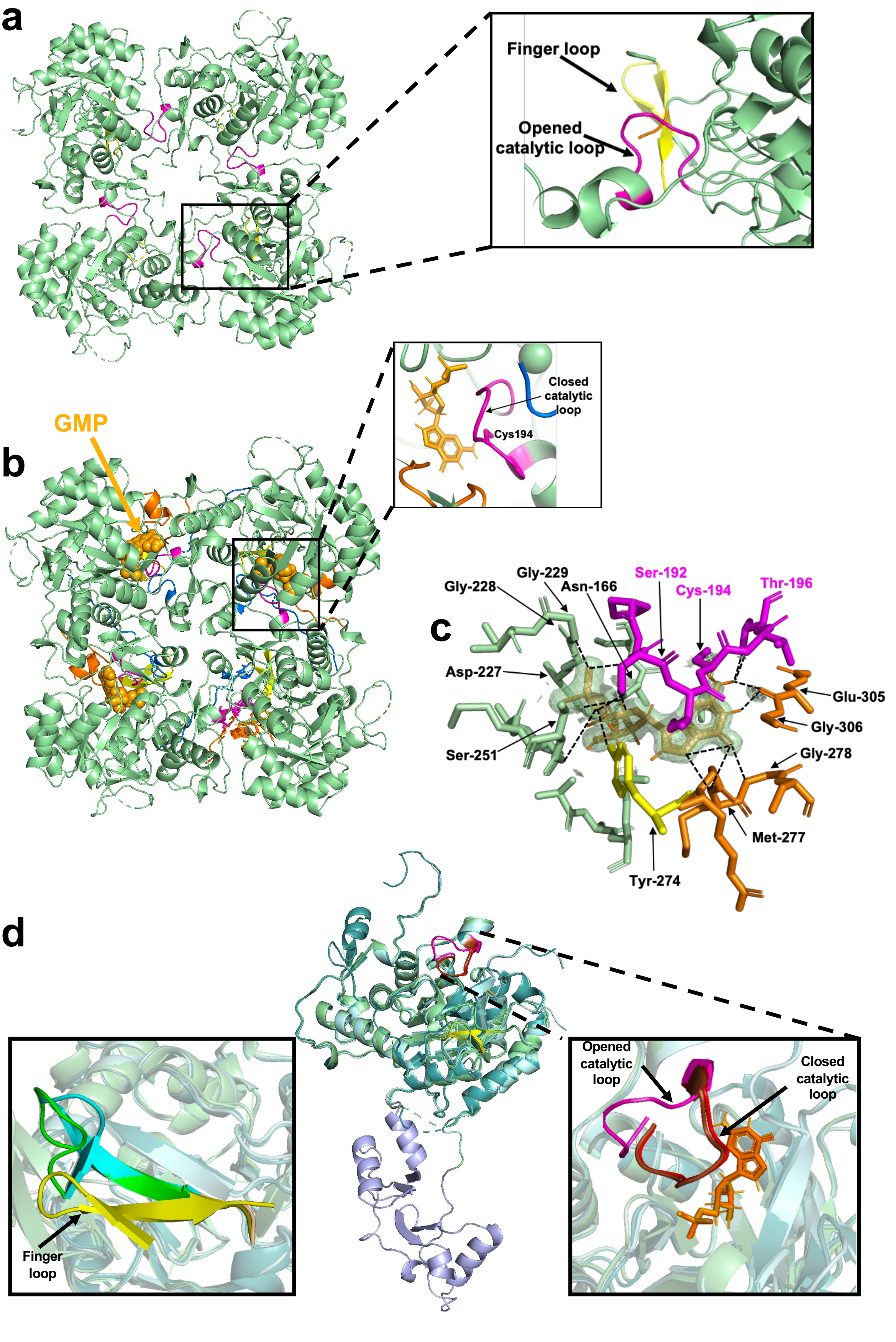

**Supplementary Figure 3- X-ray Structure of IMPDHbt ΔBD in the apo state (PDB code 9HKA) and in the presence of GMP (PDB code 9HKB). a** Ribbon representation of the tetrameric apo structure with a close-up of the catalytic and finger loops within a single monomer. **b** Same as in **a** for the complex with GMP. **c** Detailed view of the GMP-binding site in which interacting side chains are represented as light green ball-and-stick models and hydrogen bonds are shown as black dashed lines. The catalytic domain is represented in light green, GMP is shown in light orange, and key loops (catalytic, finger, flap and C-terminal) in pink, yellow, orange, and blue, respectively. **d** Comparison of the monomeric structures of IMPDHbt WT in the presence of IMP (PDB code 9HK6) with the ΔBD variants in apo (PDB code 9HKA) or GMP-bound (PDB code 9HKB) states. Binding of IMP (orange) or GMP (light orange) induces closure of the catalytic loop (red and brown) and reorientation of the finger loop (cyan and green) toward the substrate, while both loops remain (magenta and yellow, respectively) more disordered and distant from the active site in the apo state.

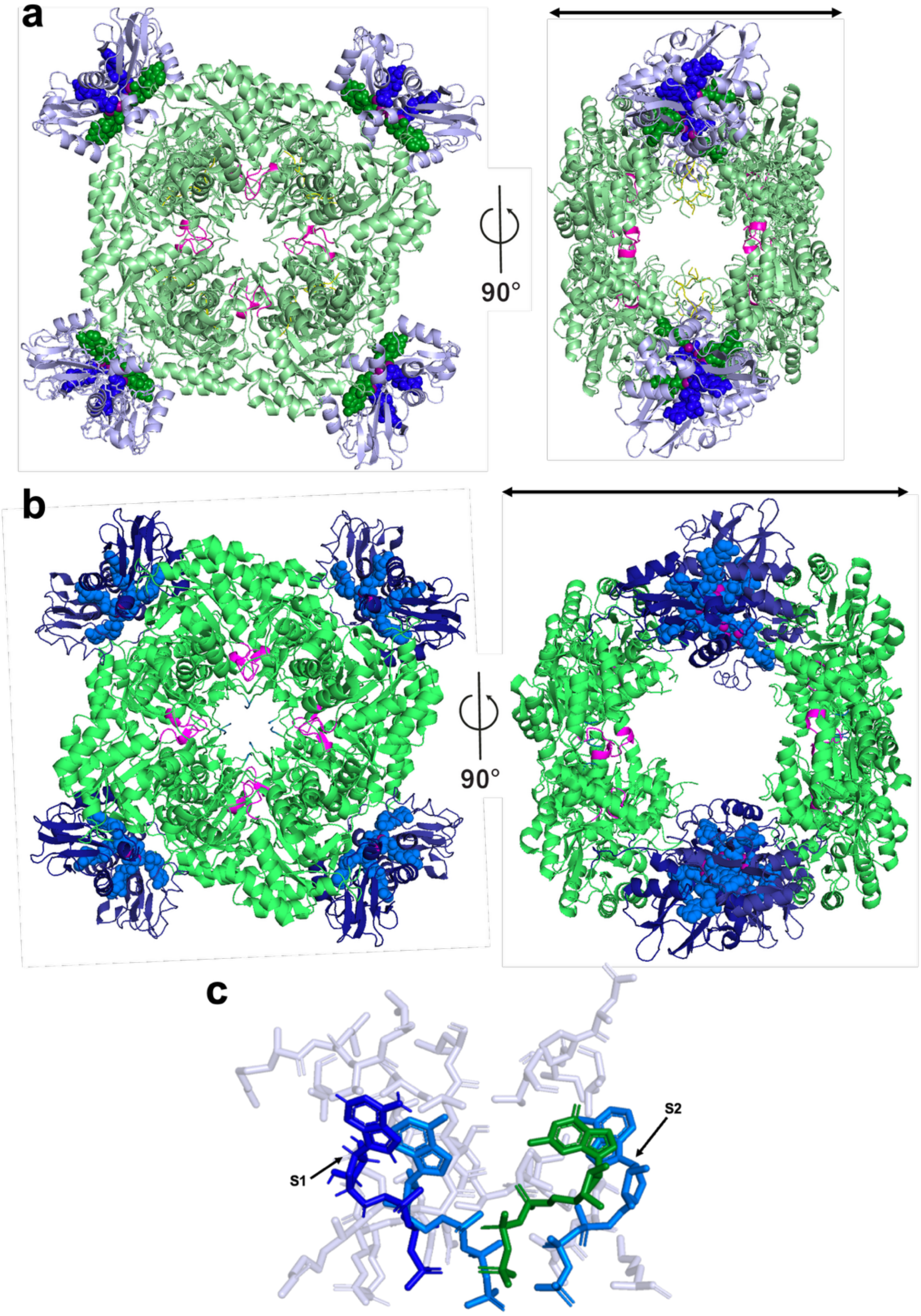

**Supplementary Fig. 4- Comparison of the crystal structures of IMPDHbt with MgATP and MgGTP (PDB code 9HK7) and IMPDHpa with MnATP (PDB code 4DQW). a-b** Ribbon representation of the octameric structures of IMPDHbt (**a**) bound to MgATP (blue) and MgGTP (green), and IMPDHpa (**b**) bound to MnATP (turquoise). Ligands are shown as spheres. **c** Superimposition of ATP and GTP ligands bound to the S1 and S2 sites in IMPDHbt and IMPDHpa. Binding sites within the Bateman domain of IMPDHbt are shown as ball-and-stick models.

**Supplementary Table 1- $IC_{50}$ values of MgGTP for IMPDHbt, IMPDHec and IMPDHpa.** $IC_{50}$ values were determined from the experimental data presented in Fig. 1c and Supplementary Fig. 1.

| | $IC_{50}$ (mM) | | |
|---|---|---|---|
| | IMPDHbt | IMPDHec | IMPDHpa |
| MgGTP | 1.41 ± 0.08 | 3.42 ± 0.65 | 0.56 ± 0.03 |

**Supplementary Table 2- Biophysical parameters derived from SAXS data (Fig. 4) of IMPDHbt.** Radius of gyration (Rg) and forward scattering intensity (I(0)) values were determined from both Guinier analysis and pair-distance distribution functions [P(r)]. Estimated molecular weights (MW) were calculated using Primus software. Relative I(0) values, which are proportional to protein concentration and molecular weight, are also reported. All measurements were performed at a protein concentration of 3 mg/mL in the presence of the indicated ligand concentration.

| | IMPDHbt | | | | | |
|---|---|---|---|---|---|---|
| | **Apo** | **IMP** 6 mM | **GMP** 5 mM | **$NAD^+$** 4 mM | **MgATP** 5 mM | **MgGTP** 5 mM |
| **I(0) Guinier ($cm^{-1}$)** | $97.07 \times 10^{-3}$ | $90.77 \times 10^{-3}$ | $88.66 \times 10^{-3}$ | $197.2 \times 10^{-3}$ | $196.3 \times 10^{-3}$ | $194 \times 10^{-3}$ |
| **Rg Guinier (Å)** | 44.99 | 44.86 | 44.54 | 52.36 | 52.43 | 49.78 |
| **I(0) P(r) ($cm^{-1}$)** | $97.07 \times 10^{-3}$ | $90.77 \times 10^{-3}$ | $88.66 \times 10^{-3}$ | $197.1 \times 10^{-3}$ | $196.3 \times 10^{-3}$ | $194 \times 10^{-3}$ |
| **Rg P(r) (Å)** | 45 | 44.9 | 44.56 | 52.27 | 52.37 | 49.73 |
| **$d_{max}$ (Å)** | 149 | 153 | 152 | 146 | 146 | 153 |
| **$MM_{Vc}$ (Da)** | 217470 | 210076 | 214055 | 475157 | 492293 | 406744 |
| **$MM_{size\&shape}$** | 206016 | 232330 | 189528 | 412100 | 533486 | 350736 |

**Supplementary Table 3. Plasmid names and composition of buffers used for purification (Buffer A) and kinetics (Buffer K) for each IMPDH (named by an acronym).** BD, Bateman domain. For size exclusion chromatography, buffer A was supplemented with 1 mM DTT and 1 mM EDTA.

<table>
<tr><th>Protein acronym</th><th>Protein description</th><th>Plasmid name</th><th>Purification buffer (Buffer A)</th><th>Kinetics buffer (Buffer K)</th></tr>
<tr><td>IMPDHbt</td><td>WT form of B. thailandensis IMPDH</td><td>pHL161-1</td><td rowspan="2">$Na_2CO_3$ 50 mM<br>pH 9,5<br>KCl 100 mM</td><td rowspan="2">$K_2HPO_4$ 50 mM<br>pH 8<br>KCl 100 mM</td></tr>
<tr><td>IMPDHbt ΔDB</td><td>BD-deleted variant of B. thailandensis IMPDH</td><td>pHL161-2</td></tr>
<tr><td>IMPDHec</td><td>WT form of E. coli IMPDH</td><td>pHL166-1</td><td>Tris-HCl 50 mM<br>pH 9<br>KCl 100 mM</td><td>Tris-HCl 50 mM<br>pH 8<br>KCl 50 mM</td></tr>
<tr><td>IMPDHpa</td><td>WT form of P. aeruginosa IMPDH</td><td>pHL143-1</td><td>$K_2HPO_4$ 50 mM<br>pH 8<br>KCl 100 mM</td><td>Tris-HCl 50 mM<br>pH 8<br>KCl 100 mM</td></tr>
</table>

**Supplementary Table 4. Summary of crystallization conditions and bound ligands for all PDB structures reported in this study.** Protein concentrations for crystallization were 8.55 mg/mL for IMPDHbt and 10 mg/mL for IMPDHbt ΔBD.
Abbreviations: PEG, polyethylene glycol; CHES, N-cyclohexyl-2-aminoethanesulfonic acid; $LiNO_3$, lithium nitrate; NaCl, sodium chloride; Na malonate, sodium malonate; MgCl2, magnesium chloride, Tris-HCl, tris(hydroxymethyl)aminomethane hydrochloride.

| Protein acronym | PDB ID | Description | Crystallization conditions | Cryo-protectant |
|---|---|---|---|---|
| IMPDHbt | **9HK6** | 5mM IMP | 10%w/v PEG 8K, 0.1 M CHES pH 9.5 | Ethylene glycol |
| | **9HK7** | 5mM MgGTP, 5 mM MgATP | 20%w/v PEG 3350, 0.2 M $LiNO_3$ | Glycerol |
| | **9HK8** | 5 mM MgGTP, 5 mM MgATP, 5 mM IMP, 4 mM NAD | 20%w/v PEG 3350, 50 mM NaCl | Glycerol |
| | **9HK9** | 5 mM MgGTP, 5mM MgATP, 5mM IMP | 20%w/v PEG 3350, 150 mM Na Malonate | Ethylene glycol |
| IMPDHbt ΔDB | **9HKA** | Apo | 16%w/v PEG 4K, 0.1 M Tris-HCl pH 8.5, 0.2 M $MgCl_2$ | Ethylene glycol |
| | **9HKB** | 5 mM GMP | 10%w/v PEG 3K, 0.1 M Na phosphate citrate pH 4.2, 0.2 M NaCl | Ethylene glycol |
